# EHD1-dependent traffic of IGF-1 receptor to the cell surface is essential for Ewing sarcoma tumorigenesis and metastasis

**DOI:** 10.1101/2023.01.15.524130

**Authors:** Sukanya Chakraborty, Aaqib M. Bhat, Insha Mushtaq, Haitao Luan, Achyuth Kalluchi, Sameer Mirza, Matthew D. Storck, Nagendra Chaturvedi, Jose Antonio Lopez- Guerrero, Antonio Llombart-Bosch, Isidro Machado, Katia Scotlandi, Jane L. Meza, Gargi Ghosal, Donald W. Coulter, M Jordan Rowley, Vimla Band, Bhopal C. Mohapatra, Hamid Band

## Abstract

Overexpression of EPS15 Homology Domain containing 1 (EHD1) has been linked to tumorigenesis but whether its core function as a regulator of intracellular traffic of cell surface receptors plays a role in oncogenesis remains unknown. We establish that EHD1 is overexpressed in Ewing sarcoma (EWS), with high EHD mRNA expression specifying shorter patient survival. ShRNA and CRISPR-knockout with mouse *Ehd1* rescue established a requirement of EHD1 for tumorigenesis and metastasis. RTK antibody arrays identified the IGF-1R as a target of EHD1 regulation in EWS. Mechanistically, we demonstrate a requirement of EHD1 for endocytic recycling and Golgi to plasma membrane traffic of IGF-1R to maintain its surface expression and downstream signaling. Conversely, EHD1 overexpression-dependent exaggerated oncogenic traits require IGF-1R expression and kinase activity. Our findings define the RTK traffic regulation as a proximal mechanism of EHD1 overexpression-dependent oncogenesis that impinges on IGF-1R in EWS, supporting the potential of IGF-1R and EHD1 co-targeting.

## 1. INTRODUCTION

Members of the EPS15 homology domain-containing (EHD) protein family (EHD1-4) of membrane-activated ATPases have emerged as key regulators of vesicular traffic along the endocytic pathway ^1, 2, 3^. Among them, EHD1 has been investigated the most and is well-established to regulate the post-endocytic recycling back to the cell surface of a variety of cell surface receptors ^1–3^. In contrast to this role in post-endocytic receptor traffic, our recent studies identified a unique role for EHD1 in the pre-activation transport of newly-synthesized RTKs, CSF1 receptor^4^ and EGFR^5^ from the Golgi to the plasma membrane to enable their efficient ligand-induced signaling and biological responses. These cell biological findings raise the possibility that overexpression of EHD1 in tumors could promote RTK-dependent oncogenic signaling by enabling the cell surface display of RTKs on tumor cells. This idea is consistent with recent findings in which EHD1 overexpression has been observed in various cancers, often correlating with shorter survival, and cell-based studies using gene knockdown or overexpression strategies that support the role of EHD1 overexpression to promote tumorigenesis, chemotherapy resistance, epithelial-mesenchymal transition, stem cell behavior and glycolysis in various tumor models ^6–15^. These studies have linked EHD1 overexpression to distal signaling alterations such as the activation of NFκB, β-catenin/c-Myc pathways that are not immediately linked to EHD1’s core vesicular traffic roles in endocytic recycling and Golgi to cell surface RTK traffic. Consistent with the potential of EHD1 expression in fact regulating RTK traffic in tumors, EHD1 levels in non-small cell lung cancer correlated with EGFR expression and specified shorter survival, metastasis, and chemotherapy resistance ^8, 16^. EHD1 was also shown to promote erlotinib resistance in EGFR-mutant lung cancers ^11^. However, direct evidence for regulation of RTK traffic as a proximal mechanism to activate the various distal signaling axes in EHD1-overexpressing cancers is currently lacking. Such a linkage is of considerable interest since receptor tyrosine kinases (RTKs) are well established as oncogenic drivers or as key secondary components of oncogenic programs of other driver oncogenes across cancers ^17^.

The oncogenesis-associated overactivity of RTKs has been ascribed to multiple mechanisms, including gene amplification, increased transcription, genetic aberrations such as chromosomal translocation, point mutations or internal deletions, alterations of downstream signaling components, as well as activation through autocrine feedback loops ^18^. A key mechanism of post-translational control of RTK levels and signaling involves the regulation of their intracellular traffic. One aspect of RTK traffic that has received the most attention is their post-activation endocytic traffic into either lysosomal degradation or the alternative recycling pathway back to the plasma membrane, with the balance of these mechanisms a key determinant of the magnitude, duration, and type of cellular responses elicited by ligand-induced RTK activation ^19^. Indeed, altered endocytic trafficking of RTKs, including the imbalance between recycling versus degradation, is now known to promote oncogenic signaling by RTKs ^20, 21^.

To investigate the potential link of EHD1 to RTK-dependent tumorigenesis, we carried out studies using Ewing Sarcoma (EWS), the second most common malignant bone tumor in children and young adults ^22^, as a model. Despite advances in multimodality treatment strategies, the EWS prognosis remains poor, with cure rates below 25%, due to its aggressive and metastatic nature ^23–25^. More than 85% of cases harbor reciprocal translocations that generate a currently undruggable fusion oncogene composed of portions of *EWS* and ETS transcription factor *FLI1* ^24^. *EWS-FLI1* drives oncogenesis through altered transcriptional activity as well as other mechanisms that together promote a fully malignant phenotype ^26, 27^.

Upregulation of signaling through multiple RTKs is implicated in EWS tumorigenesis, metastasis, and therapy resistance, with most attention to the role of insulin-like growth factor 1 receptor (IGF-1R) ^17^. IGF-1R was demonstrated to be required for EWS/FLI1-mediated transformation of EWS cells ^28^. Furthermore, EWS/FLI and other EWS-associated fusion oncoproteins transcriptionally upregulate the IGF-1 expression ^29^ EWS-FLI1 binding to IGF binding protein 3 (IGFBP-3) promoter was found to repress the expression of this key negative regulator of IGF-1R signaling, leading to constitutively active IGF-1R signaling in EWS cells ^30^. IGF-1R and components of the IGF-1 receptor signaling pathway have also been associated with the development, progression, and metastasis of breast, non-small cell lung, and other solid cancers ^31–33^. Many preclinical studies support the potential of IGF-1R targeting to limit tumorigenesis and metastasis ^32, 34, 35^. In EWS in particular, IGF-1R inhibition has been explored ^36–43^ but the results of clinical trials with antibody- and tyrosine kinase inhibitor (TKI)-based IGF-1R targeting have been disappointing ^44, 45^. The inefficacy of IGF-1R targeting in clinic likely reflects the lack of predictive markers of therapeutic response as well as our still incomplete understanding of the regulation of IGF-1R in tumors.

Given the important roles of IGF-1R and other RTKs in supporting the fusion oncoprotein-driven tumorigenesis and metastasis in EWS, we test our hypothesis that EHD1 overexpression enables high cell surface levels of RTK as a novel pro-oncogenic mechanism using EWS as a model. Our results establish a critical positive role of EHD1 overexpression in EWS oncogenesis and demonstrate that EHD1-dependent endocytic recycling and pre-activation Golgi to the plasma membrane traffic of IGF-1R are essential for its oncogenic role.

## 2. MATERIALS AND METHODS

### Ewing sarcoma patient tissue microarrays and immunohistochemical analysis

A total of 324 paraffin-embedded samples from ESFT patients from the period comprised between April 1971 and May 2007 treated at Instituto Ortopedici Rizzoli (IOR), Bologna, Italy, and at the Department of Pathology of the University of Valencia Estudi General (UVEG), Spain were analyzed within the context of two European Translational Research projects [PROTHETS (http://www.prothets.org) and EuroBo-Net (http://www.eurobonet.eu)]. All cases were genetically confirmed as belonging to the ESFT by molecular biology and/or fluorescent in situ hybridization (FISH). Approval for data acquisition and analysis was obtained from the Ethics Committee of the institutions involved in the study. The clinical data were reviewed and stored within a specific database. Characteristics of the cohort and relevant clinical information have been previously reported ^67^. A total of 24 tissue microarrays (TMAs) containing two representative cores for each case (1 mm in diameter) were constructed for immunohistochemical analysis. Out of 324 samples, 307 and 227 samples could be analyzed for EHD1 and IGF-1R IHC expression, respectively. The deparaffinized sections were stained as per standard IHC protocol. Immunoreactivity was defined as follows: negative, fewer than 5% of tumor cells stained; poorly positive (score 1), between 5% and 10% of tumor cells stained; moderately positive (score 2), between 10% and 50% of tumor cells stained, and strongly positive (score 3), with more than 50% of the tumor cells were stained.

### Cell lines and medium

Human Ewing Sarcoma cell lines TC-71, MHH-ES-1 and A4573 were obtained from Dr. Jason Yustein laboratory at Baylor college of medicine(TC-71, MHH-ES-1:DSMZ-German collection, A4573: Cellonco) and cultured in complete RPMI medium (Hyclone; #SH30027.02) with 10% fetal bovine serum (Gibco; #10437-028), 10 mM HEPES (Hyclone; #SH30237.01), 1 mM each of sodium pyruvate (Corning; #25-000-CI), nonessential amino acids (Hyclone; #SH30238.01), and L-glutamine (Gibco; #25030-081), 50 μM 2-ME (Gibco; #21985-023), and 1% penicillin/ streptomycin (#15140-122; Gibco). A673 and SK-ES-1 cells were obtained from ATCC and cultured in complete DMEM medium (Gibco; #11965-092), and complete RPMI medium supplemented as above. HEK-293T cells (ATCC CRL-3216) were cultured in complete DMEM medium. Cell lines were maintained for less than 30 days in continuous culture and were regularly tested for mycoplasma.

### Reagents and Antibodies

Primary antibodies used for immunoblotting were as follows: anti-HSC70 (#sc-7298) from Santa Cruz Biotechnology; anti-IGF-1Rβ (#3018), anti-phospho-IGF-1R-Y1135 (#3918), anti-phospho-AKT-S473 (#4060), anti-AKT (#4685), anti-ERK1/2 (#4695), anti-phospho-ERK1/2-Thr202/Tyr204 (#9101) from Cell Signaling Technology; and anti-beta-actin (#A5441) from Sigma. In-house generated Protein G-purified rabbit polyclonal anti-EHD1, EHD2, EHD3 and EHD4 antibodies have been described previously^2^. The horseradish peroxidase (HRP)-conjugated Protein A (#101023) and HRP-conjugated rabbit anti-mouse secondary antibody (#31430) for immunoblotting were from Thermo Fisher. Antibodies used for immunofluorescence studies were as follows: anti-EHD1 (#ab109311) from Abcam; Alexa-555-conjugated anti-GM130 (#48641), anti-LAMP1 (#9091) and anti-RAB11 (#5589) from Cell Signaling Technology; and anti-IGF-1Rβ (#MA5-13802) from Invitrogen. Secondary antibodies used for immunofluorescence studies were Alexa Fluor 594-conjugated goat anti-rabbit IgG (H + L) (#A11012) or Alexa Fluor 488-conjugated goat anti-mouse IgG (H + L) (#A11001) from Life Technologies Corporation. The Annexin-V-PI flow cytometric analysis was done using a kit (#V13241) from Invitrogen. Primary antibodies used for immunohistochemical studies included: anti-IGF-1R (#14534) and anti-cleaved-caspase 3 (#9661) from Cell Signaling Technology; and anti-CD99 (#ab-227738) and anti-Ki67 (#ab92353) from Abcam. For immunoprecipitation studies, primary antibodies included: anti-IGF-1Rβ (Cell Signaling technology; #9750), anti-EHD1 (Abcam; #ab109311) and anti-Rabbit-IgG (Invitrogen; #02-6102). The sources for other reagents were as follows: cycloheximide (Sigma; #C7698); bafilomycin-A1 (SelleckChem; #S1413); linsitinib (SelleckChem; #S1091); recombinant-human-IGF-1 (Peprotech; #100-11); IGF-1 Receptor α mAb(1H7) (Santa Cruz; #sc-461); doxycycline (Sigma Aldrich; #D9891); Aprotinin (Sigma Aldrich #A1153); and Leupeptin (Sigma Aldrich #L2884).

### Generation of knockdown, CRISPR knockout and luciferase reporter cell lines

To generate stable doxycycline-inducible EHD1-shRNA and non-targeting control (NTC)-shRNA expressing TC71, A673 and SK-ES-1 cell lines, the following lentiviral SMART-vector constructs encoding a GFP and human EHD1-shRNA (#V3SH11252-229594140, #V3SH11252-225446205 and #V3SH11252-228109140, designated shEHD1 #1, #2 and #3, respectively) or an NTC-shRNA were obtained from Dharmacon. Lentiviral supernatants were by transient co-transfection of individual constructs with packaging plasmids (psPAX2, Addgene #12260 and pMD2.G, Addgene #12259 into HEK-293T cells using X-tremeGENE HP DNA transfection reagent (#06366236001; Roche). Lentiviral supernatants were applied to cells for 48h in the presence of polybrene (10 µg/ml, Sigma #H9268) and stable polyclonal cell lines were selected with 1 μg/ml puromycin and maintained in their respective media with tetracycline-free 10% FBS (Novus Biologicals #S10350) and 1 μg/ml puromycin. For CRISPR-Cas9 mediated gene editing, the EHD1 sgRNA CRISPR/Cas9 All-in-One Lentivector (pLenti-U6-sgRNA-SFFV-Cas9-2A-Puro; #K0663105) or Scrambled sgRNA CRISPR/Cas9 All-in-One Lentivector (#K010) from Applied Biological Materials were used to generate lentiviral supernatants that were transduced into TC71 or A673 cell lines followed by selection with 1 μg/ml puromycin. Clonal derivatives were obtained by limiting dilution and screened for complete knockout using western blotting. Unless otherwise indicated, 3 or 4 clones (maintained separately) representing two EHD1 sgRNA targets were pooled for experimental analyses. For rescue experiments, the mouse *Ehd1* lentiviral vector (pLenti-GIII-CMV-RFP-2A-Puro) (#190510640495; Applied Biological Materials) was stably transduced into TC71-EHD1-KO, A673-EHD1-KO and SK-ES-1 cell lines followed by selection with 1 μg/ml puromycin. The tdTomato-luciferase plasmid was generated by recombineering using the following pMuLE system plasmids from Addgene: pMuLE ENTR U6-miR-30 L1-R5 (#62113); pMuLE ENTR SV40 tdTomato L5-L2 (#62157) and pMuLE Lenti Dest Luc2 (#62179). The mCherry-luciferase plasmid (pCDH-EF-eFFly-T2A-mCherry; Addgene #104833) was used to generate lentiviral supernatants that were transduced into the indicated cell lines followed by FACS sorting of mCherry-high fraction. *EHD1* knockout sites were assessed by Sanger sequencing of PCR fragments generated with genomic DNA as template with the following primers: 5’-AGTGTGGGTCGCTCCCG-3’ (forward) and 3’-GAGGAGCACCATAGGCTTGT-5’ (reverse). For IGF-1R siRNA knockdown, ON-TARGETplus SMARTpool siRNA (#L-003012-00-0005), ON-TARGETplus Non-targeting pool(#D-001810-10-05) were transiently transfected into cells using Dharmafect I transfection reagent (#T-2001-01) (all from Dharmacon – Horizon Discovery).

### Western Blotting

Whole cell extracts were prepared, and western blot was performed as described previously^5^ with minor modifications. Cells were lysed in Triton-X-100 lysis buffer (50 mM Tris pH 7.5, 150 mM NaCl, 0.5% Triton-X-100, 1 mM PMSF, 10 mM NaF, 1 mM sodium orthovanadate, 10 μg/ml each of Aprotinin and Leupeptin) Lysates were rocked at 4°C for >1 h, spun at 13,000 rpm for 30 minutes at 4°C and supernatant protein concentration determined using the BCA assay kit (#23225; Thermo Fisher Scientific). 30-50 μg aliquots of lysate proteins were resolved on sodium dodecyl sulfate-7.5% polyacrylamide gel electrophoresis (SDS-PAGE), transferred to polyvinylidene fluoride (PVDF) membrane, and immunoblotted with the indicated antibodies.

### Immunoprecipitation (IP)

1-mg aliquots of cleared lysate protein were incubated with optimized amounts of the indicated antibodies and rocked overnight at 4°C. 60 μl of PBS-pre-washed and PBS/1% BSA blocked protein A-Sepharose beads (#101042; Invitrogen) were added to each sample and rocked overnight at 4°C. The beads were washed six times with TX-100 lysis buffer, and bound proteins were resolved by SDS–7.5% PAGE, transferred to PVDF membrane, and immunoblotted with indicated primary antibodies. 50 μg aliquots of whole cell lysates were run as input controls.

### Immunofluorescence

Cells plated on Poly-L-lysine coated coverslips were treated as indicated in figure legends, fixed using 4% paraformaldehyde in PBS for 20 minutes at RT. Cells were then permeabilized in 0.3% Triton X-100 for 20 minutes at room temperature, blocked with 10% goat serum in PBS, and incubated with primary antibodies in 1% goat serum and 1% BSA in PBS at 4°C overnight. After washes in 0.1% BSA-PBS, cells were incubated with the appropriate fluorochrome-conjugated secondary antibody for 1 hour at RT, washed 0.1% BSA-PBS and mounted using Vectashield-mounting medium with DAPI (Vector Laboratories; #H-1500). Confocal images were captured using a Zeiss LSM 800 with microscope Airyscan. Merged pictures were generated using ZEN 2012 software from Carl Zeiss and fluorescence intensities were quantified using the ImageJ (NIH) software. Pearson’s correlation coefficients of Co-localization were analyzed using the ImageJ JACoP colocalization analysis module. A threshold was established first using the JACoP threshold optimizer, followed by calculation of Pearson’s correlation coefficients.

### Quantification of cell surface IGF-1R using FACS analysis

2×10^5^ cells were seeded per well of six-well plates and grown in regular medium with 10% FBS for 48 h. Cells were further treated as indicated in figure legends, rinsed with ice-cold PBS, released from dishes with trypsin-EDTA (#15400054; LifeTech (ThermoFisher)) and the trypsinization stopped by adding equal volume of soybean trypsin inhibitor (#17075029; LifeTech (ThermoFisher) Cells were washed thrice in ice-cold FACS buffer (1% BSA in PBS), and live cells stained with PE-anti-human-IGF-1R (#351806; Biolegend) or PE-Mouse-IgG isotype control (#400112; Biolegend). FACS analyses were performed on a LSRFortessa X50 instrument and data analyzed using the FlowJo software.

### Trans-well migration and invasion assay

For migration and invasion assays, 2×10^5^ cells were seeded in top chambers of regular or Matrigel-coated trans-wells (migration – Corning #353097; invasion – Corning #354480) in 400 μl of 0.5% FBS-containing medium for 3 hours before migration/invasion towards medium containing 10% FBS or 100 ng/ml IGF-1 in lower chambers, as indicated in figure legends. Both the top and lower chamber media contained Mitomycin C (10 μg/ml) to eliminate the contribution of cell proliferation. After 16 hours, the cells on the upper surface of the membranes were scraped with cotton swabs, and the migrated cells on the bottom surface were fixed and stained in 0.5% crystal violet in methanol. Five randomly selected visual fields on each insert were photographed, and cells were enumerated using the ImageJ software. Each experiment was run in triplicates and repeated three times.

### Cell proliferation assay

500 cells/well were seeded in 96-well flat-bottom plates in 100 ml medium and an equal volume of the CellTiter-Glo Luminescent Assay Reagent (#G7571; Promega) added at the indicated time-points. Luminescence was recorded using a GloMax® luminometer (Promega).

### Anchorage independent growth assay

10^4^ cells suspended in 0.4% soft agar were plated on top of a pre-solidified 0.8% soft agar bottom layer in 6-well plates. After two weeks, cells were fixed and stained with 0.5% crystal violet in methanol and imaged under a phase contrast microscope. The number of colonies in the entire well were quantified using the Image J software. All experiments were done in triplicates and repeated three times.

### Tumor-sphere assay

Cells were suspended in DMEM/F12 media (Thermo Fisher; #1133032) supplemented with 1% penicillin/streptomycin, 4 μg/ml heparin (Stem cell technologies; #07980), 20 ng/ml Animal-Free Recombinant Human EGF (Peprotech; #AF-100-15), 10 ng/ml Recombinant Human FGF-basic (Peprotech; #100-18B), 1X N-2 supplement (Gibco; #17502-048), 1X B27 supplement (Gibco; #17504-044) and 4% Matrigel (BD Biosciences; #356234) and seeded at 10^4^/well in ultra-low attachment 24-well plates in … volume. After one week, tumor-spheres were imaged under a phase contrast microscope. Tumor-spheres greater than 40 μm in diameter were quantified using the Image J software. All experiments were done in triplicates and repeated 3 times.

### RNA sequencing and enrichment analysis of differentially expressed genes

Total RNA was isolated using Qiagen RNeasy RNA extraction kit (#74104) and further cleaned using the RNeasy PowerClean Pro Cleanup kit (#13997-50), as per manufacturer’s protocols. The purity of RNA was assessed on a Bioanalyzer in the UNMC Next Generation Sequencing Facility. 1 µg of cleaned RNA samples were used to generate RNA-seq libraries using the TruSeq RNA Library Prep Kit v2 (Illumina) following the manufacturer’s protocols and sequenced using the 2 x 75 bases paired-end protocol on a NextSeq550 instrument (Illumina). For differential expression analysis, paired-end reads were aligned to the human genome version hg38 using hisat2 guided by Ensembl gene annotations^68^ and annotated transcripts were quantified and TPM normalized using Stringtie 2.1.1^69^ Differential expression was assessed by DESeq2^70^ and significantly changed genes were required to have a Benjamini–Hochberg adjusted p-value of < 0.05 and a 2-fold change in expression. Gene Set Enrichment Analysis (GSEA) and pathway analyses were performed using MSigDB and Ingenuity-Pathway Analysis (IPA).

### RNA isolation and Real Time-PCR analysis

Total RNA was extracted from cells using the Qiagen RNeasy RNA extraction kit (#74104) as per manufacturer’s protocols. cDNA was obtained by reverse transcription using the QuantiTect Reverse Transcription kit (Qiagen; #205311) and real-time qPCR was performed using the SYBR Green labeling method (Qiagen; QuantiTect SYBR Green PCR kit #204143) on an Applied Bioscience QuantStudio thermocycler. The primer sequences (Integrated DNA Technologies) for qRT-PCR were: human *IGF1R* 5’-TCTGGCTTGATTGGTCTGGC-3’(forward),5’-AACCATTGGCTGTGCAGTCA-3’(reverse); *PCNA* 5’-AGCAGAGTGGTCGTTGTCTTT-3’ (forward), 5’-TAGGTGTCGAAGCCCTCAGA-3’ (reverse); *E2F1* 5’-CGCCATCCAGGAAAAGGTGT-3’(forward), 5’-AAGCGCTTGGTGGTCAGATT-3’ (reverse); *E2F2* 5’-CAACATCCAGTGGGTAGGCA-3’(forward), 5’-TGCTCCGTGTTCATCAGCTC-3’ (reverse); *CDK4* 5’-TGTATGGGGCCGTAGGAAC-3’(forward), 5’-TCCAGTCGCCTCAGTAAAGC-3’(reverse); *CDK6* 5’-ACCCACAGAAACCATAAAGGATA-3’(forward), 5’-GCGGTTTCAGATCACGATGC-3’(reverse). The fold change of gene expression was calculated relative to the control using the ΔΔCt method and normalized to *GAPDH*.

### Phospho-RTK array analysis

The Human Phospho-RTK Array Kit from R&D systems (#ARY001B) was used. Cells grown to 80% confluency were lysed and 300 μg of lysate protein were applied to supplied arrays and processed according to manufacturer’s instructions. Signals corresponding to 49 tyrosine phosphorylated RTKs on the array were visualized using chemiluminescence and analyzed using ImageJ software; average signal (pixel density) of duplicate spots was used to calculate fold differences.

### Xenograft studies and IVIS imaging

All animal experiments were performed with the approval of the UNMC Institutional Animal Care and Use Committee (IACUC Protocol 19-017-04-FC). For analyses of EHD1-knockdown cell implants, 6-week-old female athymic nude mice (Charles River) were injected via the intratibial route with 10^6^ cells (in 100 μl cold PBS) engineered with lentiviral tdTomato-luciferase. Once palpable tumors were observed, the mice were randomly assigned into minus (-) Dox or plus (+) Dox groups (Dox at 2 mg/ml in drinking water with 1% sucrose). For analyses of EHD1-KO and mEHD1-rescued cell implants, 6-week-old male athymic nude mice were injected via the intratibial route with 2×10^5^ cells (in 20 μl cold PBS) engineered with lentiviral mCherry-enhanced luciferase. Tumor growth was monitored biweekly for up to 30 days using calipers, with tumor volume calculated from length x width^2^/2. For bioluminescent imaging, mice received an intraperitoneal injection of 200 μl D-luciferin (15 mg/ml; Millipore Sigma #L9504) 15 min before isoflurane anesthesia and were placed dorso-ventrally in the IVIS™ Imaging System (IVIS 2000). Images were acquired using the IVIS Spectrum CT and analyzed using the Living Image 4.4 software (PerkinElmer). Mice were imaged weekly and followed for up to 30 days. At the end of the study, mice were euthanized, and hind limbs, lungs and livers harvested. Bioluminescent signals from the harvested lungs and livers were recorded for analyses of tumor metastasis. Resected tumor xenografts were fixed in formalin, and paraffin-embedded tissue sections were used to perform the immunohistochemical staining.

### Bone quality analysis by micro-CT

The hind legs of mice harvested post-euthanasia were fixed in formalin and scanned using a micro-CT instrument (Skyscan 1172, Bruker). The parameters were 55 kV, 181 μA, 0.5 mm aluminum filter, 9 μm resolution, 4 frames averaging, 0.4 rotation step, 180° scanning. The raw images were reconstructed using the NRecon software (version 1.7.4.6, Bruker microCT). All reconstructed images were registered and realigned before analysis using the DataViewer software (version 1.5.6.2, Bruker microCT). The tibial bone was then evaluated using CTAn software (version 1.18.8.0, Bruker microCT) to calculate the percent bone volume (BV/TV), trabecular thickness (Tb.Th), trabecular number (Tb.N) and trabecular separation (Tb.Sp).

### Statistical analysis

GraphPad Prism software (version 8.0.2) was employed to perform all the statistical analyses. Statistical analyses of *in vitro* data were performed by comparing two groups using two-tailed student’s t test. Two-way ANOVA test was used to analyze the *in vivo* mouse tumor growth. P values equal to or <0.05 were considered significant. For patient tissue sample analyses, association with categorical histopathological parameters was assessed using a chi-square test to determine homogeneity or linear trend for ordinal variables. The significance level was set at 5%. To study the impact of the histological, immunohistochemical and molecular factors on progression-free survival (PFS) and disease-specific survival (DSS), the Kaplan-Meier proportional risk test (log rank) was used.

## 3. RESULTS

### EHD1 is overexpressed in EWS patient tumors and correlates with shorter event-free and overall survival

To assess if EHD1 is overexpressed in EWS patient tumors and if its overexpression bears any relationship with patient survival, we queried the publicly-available EWS patient tumor mRNA expression data using the R2 Genomics Analysis and Visualization Platform. Dichotomization of EHD1 mRNA expression levels into EHD1-High and EHD1-Low groups (mRNA expression cutoff: 439.8 TPM for event-free and 490.8 TPM for overall survival) followed by Kaplan-Meier survival analysis revealed that high EHD1 mRNA overexpression correlated with shorter event-free and overall survival in EWS patients (Fig. 1A-B). To assess if EHD1 expression is detectable in EWS patient tumors at the protein level, we carried out an immunohistochemistry (IHC) analysis of EHD1 expression in a tissue microarray of 324 EWS patient tumors. 88.6% of the 307 evaluable samples showed high EHD1 expression (IHC staining intensity of 2 or 3), while 7.49% showed low EHD1 expression (staining intensity of 1) with 3.91% deemed as negative (staining intensity of 0) (Fig. 1C-D). The level of EHD1 expression was significantly higher in metastases vs. the primary tumors (Fig. 1E). While limited survival data disallowed survival analyses, the IHC data further supported the idea that high EHD1 expression is a feature of a majority of EWS patient tumors. Overall, these analyses supported a potential pro-oncogenic role of EHD1 in EWS.

**Figure 1.**
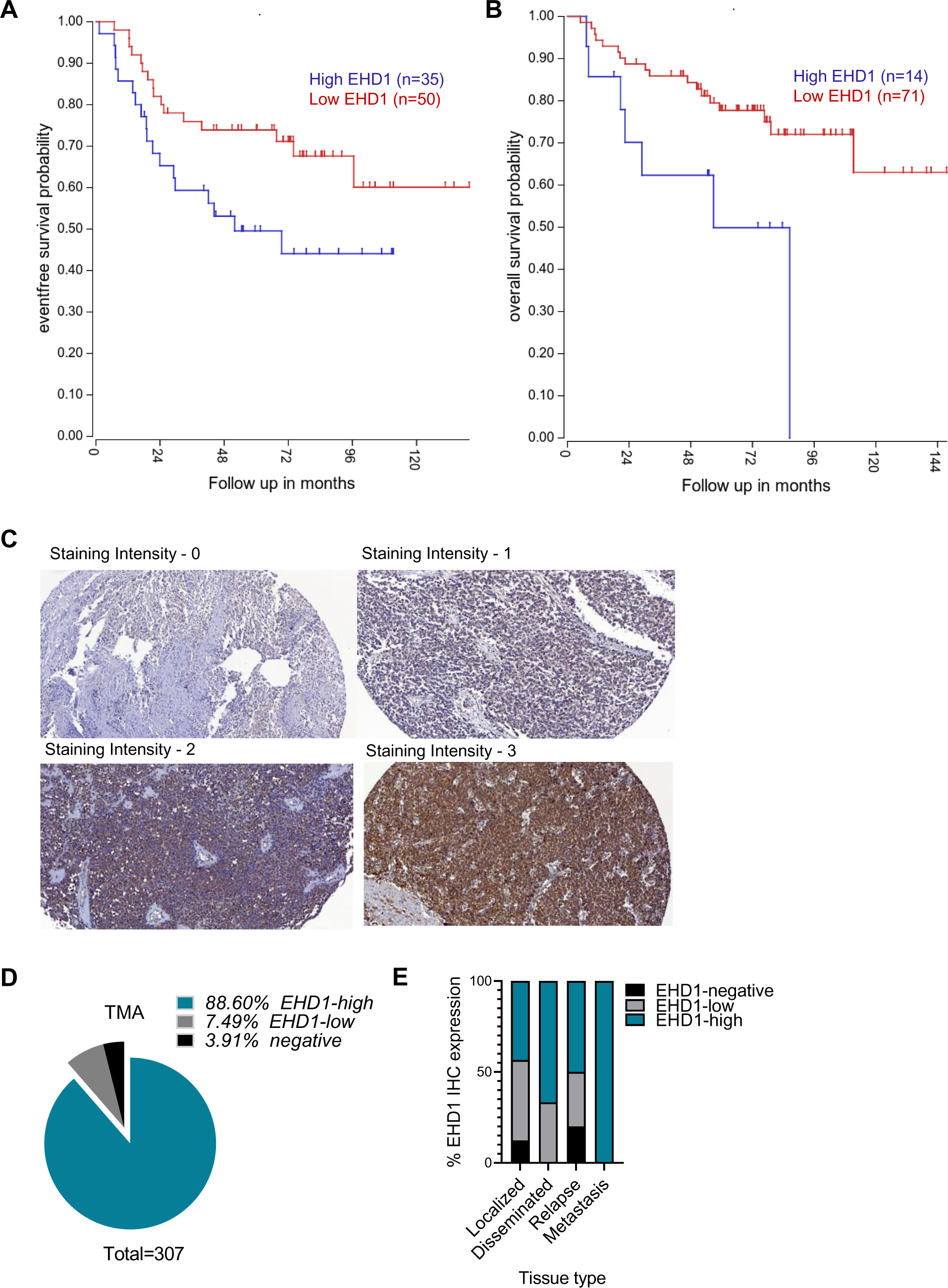
EHD1 is overexpressed in Ewing Sarcoma patient tumors and its overexpression is associated with shorter survival. (A-B) Kaplan-Meier survival analysis of 85 EWS patients based on publicly-available EHD1 mRNA expression using the R2 Genomics Analysis and Visualization Platform. EHD1-high (blue); EHD1-low (red). Event-free survival analysis (**A**; p=0.038) used a dichotomization cut-off of 439.8 (right panel), with N=35 for EHD-high and N=50 for EHD1-low group. Overall survival analysis (**B;** p=0.014) used a dichotomization cut-off of 490.8 (right panel) with N=14 for EHD1-high and N=71 for EHD1-low groups. The dichotomization cutoffs represent the program-selected defaults based on statistical significance. **(C-D)** EHD1 overexpression in EWS patient primary tumor tissue microarrays examined by immunohistochemistry (IHC). **(C)** Representative examples of various intensities (on a scale of 0 to 3) of anti-EHD1 antibody staining; details in Methods. **(D)** Relative distribution of EHD1-high (staining intensity of 2 or 3), EHD1-low (staining intensity of 1) or EHD1-negative samples (N= 307). **(E)** Significantly higher expression of EHD1 in metastatic lesions as compared to localized disease, χ^2^= 22.389; p = 0.001, Spearman’s correlation coefficient= 0.211; p < 0.001

### EHD1 is required for the maintenance of *in vitro* pro-tumorigenic and pro-metastatic oncogenic traits of EWS cell lines

To identify EWS cell models suitable for delineating the role of EHD1 in tumor biology, we first queried the CCLE database and found that most of the 19 included EWS cell lines expressed moderate EHD1 mRNA levels relative to the total cell line panel (Supplementary Table 1). Analysis of a subset of EWS cell lines representing the three EWS-FLI1 fusion oncogene types (TC71 and A673 -Type I EWS-FLI1 fusion; MHH-ES1, SK-ES-1 - Type II EWS-FLI1 fusion; and A4573 - Type III fusion) by immunoblotting revealed a good correlation between mRNA and protein levels, with consistently lower EHD1 protein levels in SK-ES-1 compared to the other 4 EWS cell lines, which showed robust EHD1 expression (A4573 is absent in the CCLE data) (Supplementary Fig S1A). While EHD2 was undetectable, all cell lines showed EHD3 expression with variable levels of EHD4. Based on these results, we used lentiviral constructs to engineer TC71, A673, and SK-ES-1 cell lines stably expressing 3 distinct doxycycline (Dox)-inducible EHD1-specific shRNAs (shEHD1) or a non-targeting control shRNA (shNTC). The shEHD1 #2 and #3 lines with robust EHD1 knockdown (KD), specifically upon Dox treatment (Fig. 2A), were selected for further analyses.

**Figure 2.**
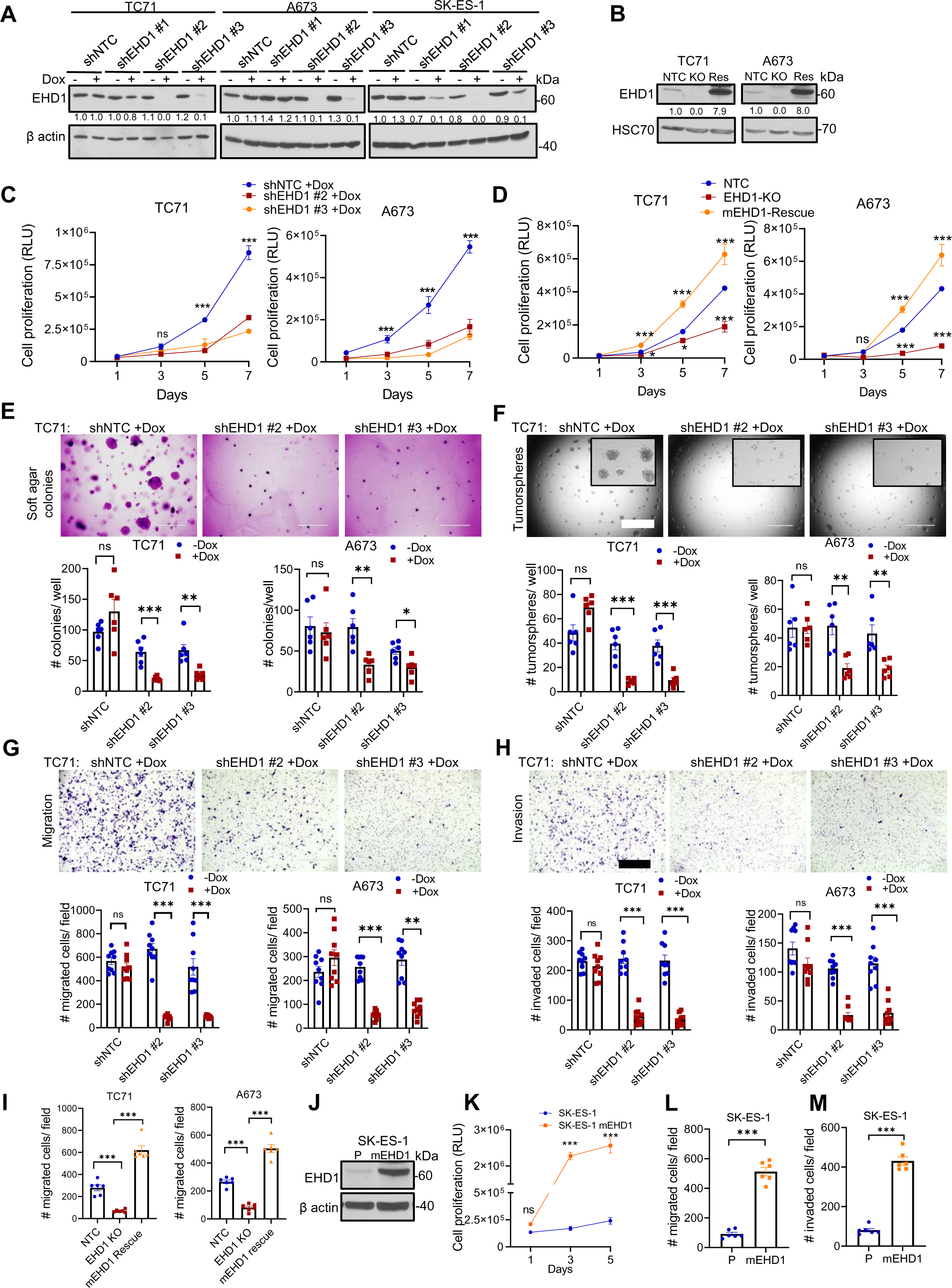
EHD1 is required to sustain the *in vitro* oncogenic traits of Ewing Sarcoma cell lines. (A) Western blot analysis of Doxycycline (Dox)-inducible knockdown of EHD1 in the indicated EWS cell lines. Cells stably expressing the non-targeting control shRNA (shNTC) or EHD1-specific shRNAs (shEHD1 #1, 2 or 3) were grown for 72h without (-) or with (+) 0.5 µg/ml Dox before lysis and immunoblotting. β-actin served as a loading control. **(B)** Western blot analysis of CRISPR-Cas9 based EHD1-KO in EWS cell lines and their derivatives with mouse EHD1 (mEHD1) expression. The indicated cell lines engineered with non-targeting control (NTC) or EHD1-targeted Cas9-sgRNA (KO) two-in-one constructs or the KO lines with mEHD1 rescue (Res) were analyzed for EHD1 expression, with β-actin served as a loading control. **(C)** Impaired cell proliferation upon EHD1 knockdown. The indicated shNTC and shEHD1 TC71 or A673 cell lines pre-treated with Dox for 48 h were plated in 96-well plates and cell proliferation assessed at the indicated time points using the Cell-Titer-Glo assay. Y-axis, Relative Luminescence Units (RLU) as a measure of increase in the number of viable cells. Data points represent mean +/− SEM of three experiments, each with six replicates. **(D)** Impaired cell proliferation upon EHD1 knockout (KO) and rescue of proliferation defect by mEHD1. Cell proliferation was assessed as in C. **(E)** Impaired soft agar colony formation upon EHD1 knockdown. The indicated shNTC and shEHD1 TC71 or A673 cell lines pre-treated with Dox for 48h were plated in soft agar and the colony numbers quantified after 3 weeks of culture in the presence of Dox. Top, representative images of TC71 cells; scale bar, 1000 μm. Bottom, mean +/− SEM of two experiments each in triplicates. **(F)** Impaired tumor-sphere formation upon EHD1 knockdown. Top, Representative images of TC71 cells; scale bar, 1000 μm. Bottom, Mean +/− SEM of two experiments each in triplicates. **(G)** Impaired trans-well cell migration upon EHD1 knockdown. Top, representative images of TC71 cells; scale bar, 400 μm. Bottom, quantification of the number of migrated cells per high-power field; mean +/− SEM of three experiments each in triplicates. **(H)** Impaired invasion through Matrigel-coated trans-wells upon EHD1 knockdown. Top, Representative images of TC71 cells; scale bar, 400 μm. Bottom, quantification of the number of invaded cells per high-power field; Mean +/− SEM of three experiments each in triplicates. **(I)** Impaired trans-well cell migration upon EHD1 knockout (KO) and rescue of migration defect by mEHD1. Analyses done as in G. **(J)** Immunoblot analysis demonstrating mEHD1 overexpression relative to endogenous EHD1 in parental cells (P) in SK-ES-1 cells **(K)** Increased cell proliferation of SK-ES-1 cells upon mouse EHD1 (mEHD1) overexpression by Cell-Titer-Glo assay. Data points represent mean +/− SEM of 3 experiments each with six replicates. **(L-M)** Transwell migration and invasion assays in SKES1-mEHD1 cells as compared to control cells. Data points represent mean +/− SEM of two experiments each in triplicates; *p<0.05, **p<0.01, ***p<0.001, ns= not significant.

First, we examined the impact of Dox-induced EHD1 KD on the various *in vitro* oncogenic traits. EHD1-KD markedly and significantly reduced the magnitude of cell proliferation, measured using the Cell-Titer Glo assay in TC71, A673, and SK-ES-1 cell lines (Fig. 2C, Supplementary Fig S1B). Furthermore, EHD1-KD in A673 and TC71 cell lines induced a significant reduction in anchorage-independent growth on soft-agar and tumor-sphere forming ability (Fig. 2E-F, Supplementary Fig. S1C-D). EHD1-KD also induced a drastic reduction of trans-well cell migration and invasiveness (migration through Matrigel) (Fig. 2G-H and Supplementary Fig. S1E-H). Treatment with the cell-proliferation inhibitor mitomycin-C excluded the role of reduced cell proliferation as a major contributor to reduction in migration and invasion; the modest reduction in proliferation in 24 hours could not account for the nearly 85% reduction in migration and invasion ability.

To further establish the pro-oncogenic role of EHD1 and its specificity, we generated CRISPR-Cas9 *EHD1* knockout (KO) derivatives of TC71 and A673 cell lines and then used a lentiviral construct to stably express mouse *Ehd1* (mEHD1) in the EHD1-KO cell lines to assess the rescue of any functional deficits (Fig. 2B, Supplementary Fig. S1K). Indeed, EHD1-KO induced a pronounced decrease in the cell proliferation and migratory ability, and this deficit was rescued mEHD1 (Fig. 2D,2I, Supplementary Fig. S1I); consistent with higher levels of the introduced mEHD1, the rescued cell lines displayed increased proliferation and migration relative to parental lines. Further illustrating the pro-oncogenic role of EHD1 overexpression, introduction of mouse *Ehd1* into EHD1-low SK-ES-1 cell line led to a marked and significant increase in cell proliferation, migration and invasion compared to parental cells (Fig 2J-M, Supplementary Fig. S1J). RNA-seq analysis showed a marked reduction in cell cycle regulatory gene expression Dox-treated shNTC vs. shEHD1 EWS cell lines among the significantly downregulated pathways (Supplementary Fig. S2A-F) and qPCR analysis validated the downregulation of *CDK4*, *CDK6*, *E2F1*, *E2F2*, and *PCNA* mRNA levels (Supplementary Fig. S2D). Collectively, our KD, KO, rescue, and overexpression analyses strongly support a key positive role of EHD1 in promoting pro-tumorigenic and pro-metastatic traits of EWS cells.

### EHD1 is required for *in vivo* EWS tumorigenesis

To assess if the marked reduction in pro-oncogenic traits seen *in vitro* translates into impaired tumorigenesis *in vivo,* we implanted TC71 NTC, EHD1-KO, and mEHD1 rescue cell lines engineered with a lentiviral mCherry-enhanced luciferase reporter ^46^ in the tibias of Nude mice (n=8 per group at the beginning) and monitored the tumor growth by luminescence imaging. While the NTC tumors exhibited time-dependent growth (seen as an increase in log10 photon flux), the EHD1-KO tumors failed to grow and, in fact, showed a reduction in photon flux; in contrast, implants of mEHD1-rescued EHD1-KO cells exhibited rapid tumor growth with higher photon flux, and mice in this group reached the euthanasia endpoints a week earlier (Fig. 3A-B, Supplementary Fig S3A-B). IVIS imaging of lungs resected at necropsy revealed detectable metastatic seeding in 3 of 8 mice implanted with NTC cells but not in any of the mice implanted with EHD1-KO cells. In contrast, 7/8 mice implanted with mEHD1-rescued cells showed metastases (Fig. 3C-D). Notably, 1/8 NTC and 2/8 rescued cell line-implanted mice exhibited liver metastases. Morphometric analysis of tibial bone by micro-CT scanning showed reduced bone volume, trabecular number, thickness, and separation in mice implanted with NTC or mEHD1-rescued TC71 cells, indicative of increased tumor-induced bone degradation, with a significant amelioration of these defects in tibias of mice implanted with EHD1-KO cells (Fig. 3E-F).

**Figure 3.**
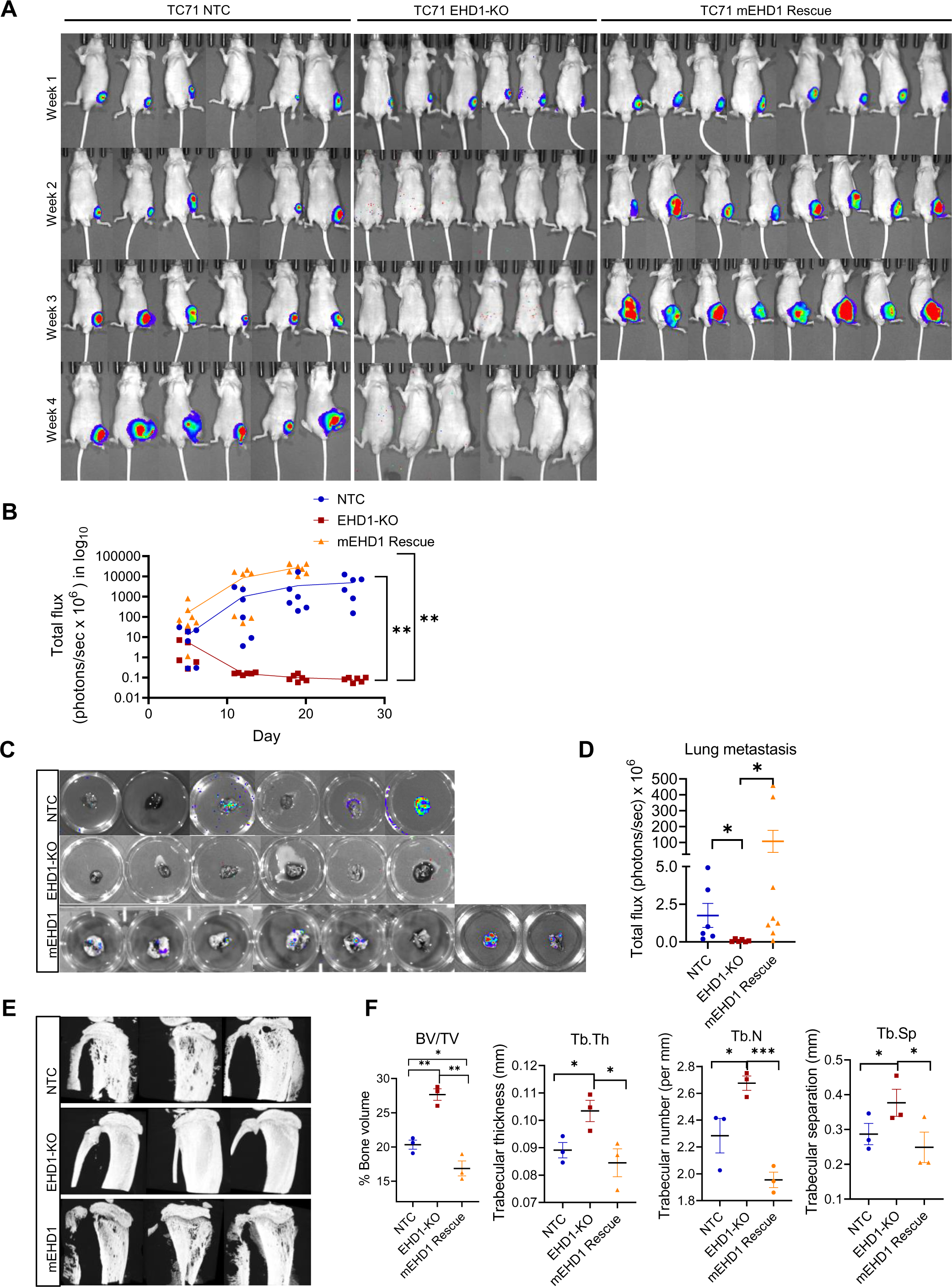
Loss of EHD1 expression markedly impairs the growth and metastasis to lungs of bone-implanted EWS cells. 2 × 10^5^ TC71 cells edited with non-targeting (NTC) or EHD1-targeted sgRNA (EHD1-KO), or the EHD1-KO cells rescued with mEHD1, all carrying a mCherry-luciferase reporter were injected in tibias of 6-week-old nude mice (8/group) and primary tumor growth was monitored by bioluminescence imaging at the indicated time points in mice with detectable bioluminescent signals at the outset (6 for NTC and EHD1-KO groups; 8 for Rescue group). **(A)** Images of individual mice with super-imposed luminescence signals over time. **(B)** Plots of log total flux values over time. Differences between groups were analyzed using two-way ANOVA; **p<0.01. **(C-D)** Bioluminescence signals of lungs harvested at necropsy are shown as individual images (C) and as quantified log total flux (D). **(E)** Micro-CT scanned images of tibias isolated from mice in the indicated groups. 3 mice per group were scanned. **(F)** Quantification of percent bone volume (BV/TV), trabecular thickness (Tb.Th), trabecular number (Tb.N) and trabecular separation (Tb.Sp) of scanned images from E, by CTAn software. Data represent Mean +/− SEM, (*p<0.05, **p<0.01, ***p<0.001).

To assess the impact of inducible EHD1 KD on pre-formed tumors, we implanted Nude mice with shNTC or shEHD1 (#3) TC71 cell lines carrying the TdTomato-luciferase reporter and monitored the tumor growth by IVIS imaging, as above. Groups of tumor-implanted mice (n=7/group for NTC and 6/group for shEHD1) were either followed as such or switched to Dox-containing water from Day 10. Comparable time-dependent growth of shNTC TC71 implants without or with Dox treatment excluded any impact of Dox itself; in contrast, the growth of shEHD1 TC71 tumors was markedly reduced by Dox treatment compared to untreated mice (p<0.0001; Supplementary FigS4A-B). Western blotting of resected tumor lysates confirmed the Dox-induced EHD1 KD in the shEHD1 group, and IHC staining with anti-human CD99 confirmed the tumor mass (Supplementary Fig.S4C-D). Tumors of Dox-treated shEHD1-implanted mice showed fewer proliferating tumor cells (Ki-67 staining) and an increase in apoptotic cells (cleaved-caspase3) (Supplementary FigS2E). Collectively, these results unequivocally demonstrate a requirement of EHD1 for EWS tumorigenesis and metastasis.

### Identification of IGF-1R as an EHD1 target in EWS

Given our prior identification of EHD1 as a regulator of Golgi to plasma membrane traffic and subsequent signaling of EGFR and CSF-1R ^4, 5^, we hypothesized that regulation of RTKs may underlie the requirement of EHD1 in EWS oncogenesis. We, therefore, probed a phospho-RTK profiling array incorporating 49 of 58 human RTKs with lysates of untreated (control) vs. Dox-treated (KD) shEHD1 #3 TC71 or A673 cell lines. The levels of phospho-IGF-1R were specifically reduced upon Dox treatment of both cell lines, while changes in other phospho-RTKs were not seen in both (Fig. 4A). Consistent with our findings with EGFR and CSF1R ^5^, analysis of TC71 cell lines harboring two distinct shRNAs (#2 or #3) demonstrated a reduction in total IGF-1R levels upon EHD1-KD (Fig. 4B). These results were further validated using control vs. CRISPR-KO TC71 and A673 cell lines; notably, mEHD1-rescued KO cell lines exhibited higher total IGF-1R levels than the non-targeted controls, consistent with higher mEHD1 levels compared to that of endogenous EHD1 in control cells (Fig. 4C). qPCR analyses demonstrated comparable IGF-1R mRNA levels between the NTC and EHD1-KO cell lines, excluding EHD1 regulation of IGF-1R levels at the mRNA level (Supplementary Fig. S5A).

**Figure 4.**
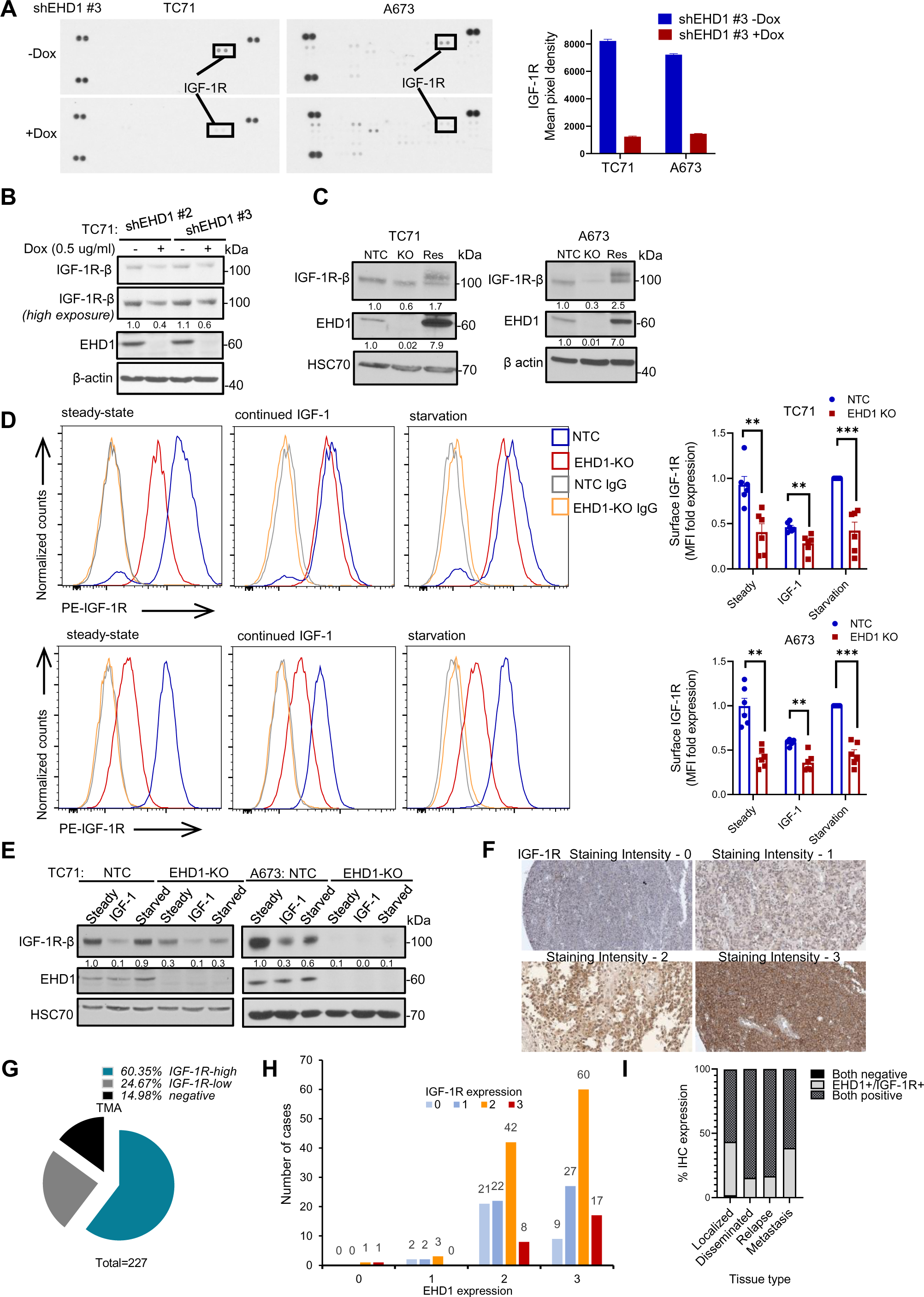
Identification of insulin like growth factor-1 receptor (IGF-1R) as a regulatory target of EHD1 in EWS. (A) Phospho-RTK antibody array analysis. Membranes arrayed with antibodies against phosphorylated versions of 49 human RTKs (each in duplicate) were probed with lysates of TC71-shEHD1 or A673-shEHD1 treated with or without Dox. Left, Images of membranes with IGF-1R spots indicated. Right, Densitometric quantification of IGF-1R signals. **(B)** Western blot showing reduced total IGF-1R protein levels in TC71 cells upon Dox-induced EHD1 knockdown. Cell lysates were probed with an anti-IGF-1Rβ antibody with β actin as a loading control. **(C)** Reduction in total IGF-1R levels upon EHD1-KO and rescue by mEHD1 expression in EHD1-KO EWS cells. Lysates of the indicated cell lines probed with anti-IGF-1Rβ antibody; HSC70 or β actin served as loading controls. **(D)** EHD1-KO leads to reduced cell surface expression of IGF-1R on EWS cell lines. Control and EHD1-KO TC71 (top panel) and A673 (bottom panel) cells were grown in regular medium (steady-state), stimulated with IGF-1 (100 ng/ml) for 16 hours prior to analysis to promote the IGF-1R degradation (continued IGF-1), or cells pre-treated with IGF-1 were switched to low serum-containing and IGF-1-free medium (starvation) to promote the cell surface accumulation of newly-synthesized and recycled IGF-1R. Live cells were stained with anti-IGF-1R or IgG control antibody and analyzed by FACS. Left, representative histograms. Right, quantification of surface IGF-1R expression. Data represents the fold ratio of Median fluorescence intensity (MFI) relative to NTC cells under starvation condition (assigned a normalized value of 1). mean +/− SEM of six independent experiments. N=3, (*p<0.05, **p<0.01, ***p<0.001, ns= not significant). **(E)** Representative immunoblotting (with densitometric quantification) for total IGF-1R expression in samples analyzed under D. (**F-I)** Positive correlation of EHD1 and IGF-1R expression in EWS patient tumors. Anti-IGF-1R IHC staining was carried out on TMAs from the same patient cohort as that analyzed for EHD1 expression (in Fig. 1). **F** shows the representative examples of the IGF-1R staining intensity of 0-3 **G** shows the relative distribution of high (staining intensity of 2-3; 60.35%), low (staining intensity of 1; 24.67%) or negative (staining intensity of 0; 14.9%) IGF-1R staining among 227 evaluable patients. **H** shows the correlation between EHD1 and IGF-1R staining intensities. Y-axis, number of cases displaying IGF-1R staining intensities of 0,1, 2 or 3. X-axis, EHD1 staining intensities, 0-3. Spearman’s Correlation Coefficient= 0.179, p=0.009. **I** shows expression of EHD1 and IGF-1R in localized disease, disseminated, relapse and metastatic lesions.

The cell surface levels of RTKs determine their access to ligands and hence the downstream responses ^20^. To assess if EHD1 is required for cell surface IGF-1R expression, we carried out live-cell IGF-1R immunostaining followed by FACS analysis on control vs. EHD1-KO TC71 and A673 cell lines under three distinct conditions: 1. Cells cultured in regular medium with 10% FBS (steady state). 2. Cells in regular medium treated with IGF-1 (100 ng/ml) to promote ligand-induced internalization and degradation of IGF-1R. 3. Cells in regular medium treated with IGF-1 (100 ng/ml) for 16 hours to promote the downregulation of cell surface IGF-1R followed by culture in low serum (0.5%) medium without added IGF-1 for 24 hours to allow the newly-synthesized receptor to accumulate at the cell surface. The cell surface IGF-1R on control cells decreased upon IGF-1 treatment followed by an increase when cultured in low-serum/IGF-1-free medium, reflecting the transport of newly synthesized IGF-1R to the cell surface (Fig. 4D). The EHD1-KO cells, in contrast, exhibited lower cell surface levels under all conditions, and the extent of IGF-1-induced surface IGF-1R downregulation was smaller than in control cells (Fig. 4D). Concurrent immunoblotting confirmed the lower IGF-1R levels in KO cells under all conditions examined (Fig. 4E). Immunofluorescence microscopy further confirmed the lower cell surface IGF1R levels in EHD1-KO compared to control TC71 or A673 cell lines (Supplementary Fig. S5B). Notably, anti-IGF-1R IHC of the EWS patient TMAs (same as those used for EHD1 staining) showed that 60.35% of the 227 interpretable samples exhibited high (staining intensity of 2-3) IGF-1R staining (Fig. 4F-G), with a positive correlation (Spearman’s Correlation Coefficient = 0.179) between EHD1 and IGF-1R staining (Fig. 4H-I).

### EHD1 controls the cell surface levels of IGF-1R by regulating its intracellular traffic itinerary

EHD1 is known to facilitate the recycling of many non-RTK receptors following their endocytosis via the Rab11+ endocytic recycling compartment ^47^ but whether EHD1 regulates RTK recycling, a key mechanism to counteract the alternate lysosomal delivery and degradation after ligand-induced internalization ^20^, is unknown. Consistent with EHD1-dependent RTK recycling, we previously observed that EHD1 colocalizes with an oncogenic kinase-active mutant or wildtype EGFR in endocytic compartments ^5^. Furthermore, ectopically overexpressed IGF-1R and EHD1 were shown to co-immunoprecipitate (co-IP), partially in an IGF-1 dependent manner, and to colocalize in intracellular vesicular compartments post-IGF-1 stimulation ^48^.

To test the role of EHD1 in regulating the itinerary of pre-existing cell surface IGF-1R, we first carried out co-IP analyses of endogenous IGF-1R and EHD1 in lysates of TC71 and A673 cells that were serum/IGF-1-deprived for 24h and then left unstimulated or stimulated with IGF-1 (50 ng/ml) for 1h. EHD1/IGF-1R complexes were seen both under unstimulated and IGF-stimulated conditions (Fig. 5A). Confocal imaging demonstrated that most IGF-1R was localized at the cell surface post-starvation, with a small intracellular pool colocalizing with EHD1; upon IGF-1 stimulation, a significantly larger intracellular, presumably endosome-localized, pool of IGF-1R colocalized with EHD1 (Supplementary Fig. S6A-B). To assess if the intracellular colocalization of EHD1-IGF-1R reflects a role of EHD1 in endocytic recycling of cell surface IGF-1R, serum/IGF-deprived (starved) control or EHD1-KO EWS cell lines were treated with cycloheximide (CHX) to inhibit further protein synthesis and pulsed with IGF-1 to promote IGF-1R endocytosis followed by chase in IGF-1-free medium for various times. Confocal imaging demonstrated that internalized IGF-1R became colocalized with the endocytic recycling compartment marker RAB11 in control cells (0 min chase) but subsequently (30- and 60-min chase) reappeared at the cell surface with a decrease in the RAB11-colocalizing intracellular signal, indicating efficient recycling; in contrast, EHD1-KO cells, showed continued IGF-1R/RAB11 colocalization during chase with lower cell surface levels. (Fig. 5B, Supplementary Fig S7A). These results support the role of EHD1-dependent endocytic recycling as one mechanism by which it sustains the cell surface levels of IGF-1R.

**Figure 5.**
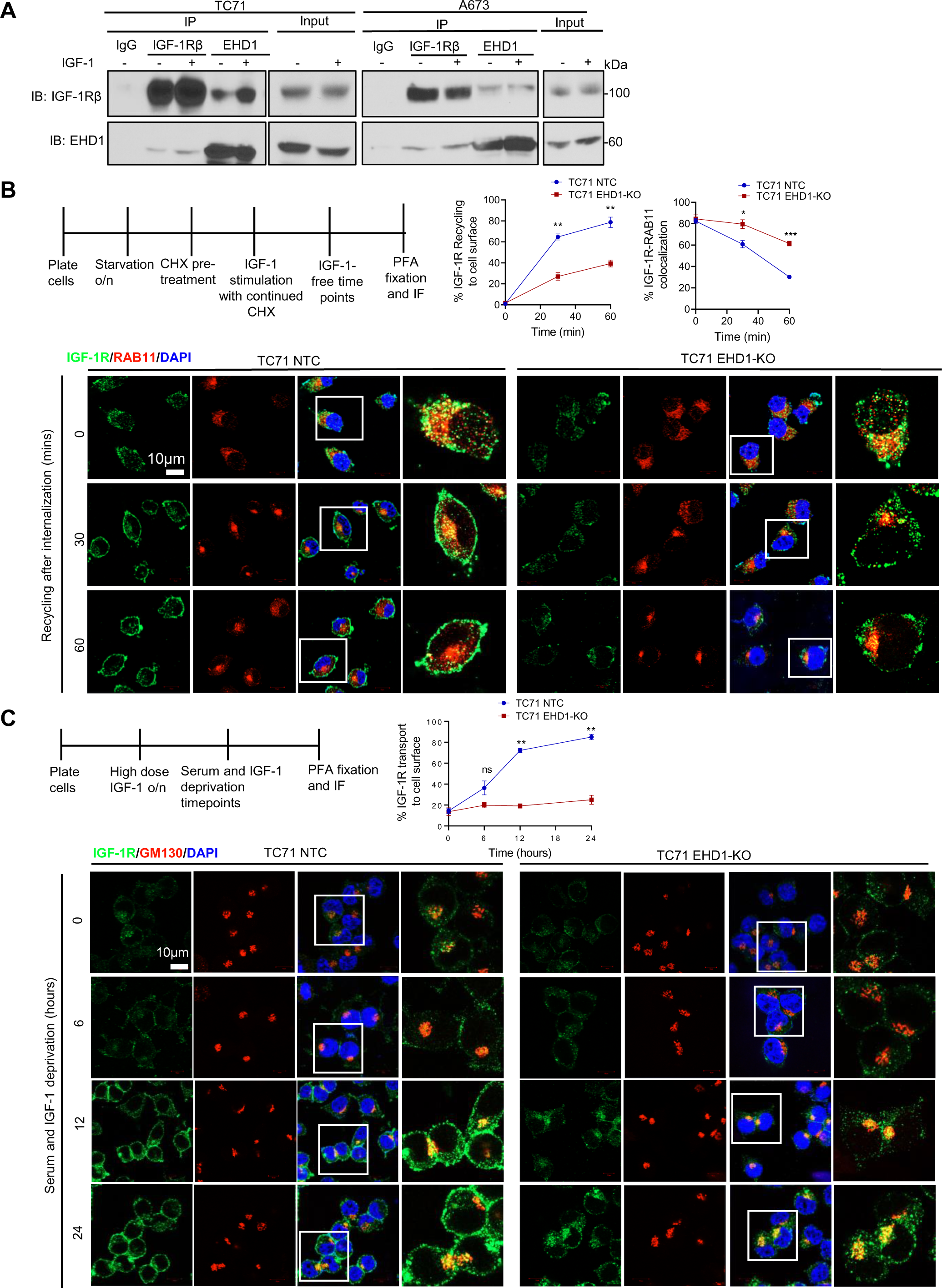
EHD1 controls cell surface IGF-1R levels by regulating its endocytic recycling and Golgi to the plasma membrane traffic. (A) EHD1-IGF-1R association in EWS cells. Anti-IGF-1Rβ or anti-EHD1 antibody immunoprecipitates (IP) from 1 mg lysate protein aliquots of the indicated cell lines were subjected to Western blotting for IGF-1Rβ or EHD1; co-IP is observed in both directions. **(B)** EHD1-KO impairs IGF-1R endocytic recycling. TC71 NTC or EHD1-KO cells pretreated with cycloheximide (50 µg/ml) for 2h to prevent new protein synthesis were treated with IGF-1 to promote the ligand-induced IGF-1R internalization (time 0), followed by incubation in IGF-1-free medium (30 and 60 min). Fixed and permeabilized cells were co-stained for IGF-1Rβ (green), RAB11 (recycling endosome marker; red) and nuclei (DAPI, blue), and analyzed using confocal imaging to assess the delivery of IGF-1R into recycling endosomes and its subsequent recycling to the cell surface. Top left, a schematic of the treatments. Bottom, Co-staining for IGF-1R and RAB11. The zoomed in panels (4^th^ columns for each cell line) show high co-localization of IGF-1R and Rab11+ in TC71-NTC cells at time 0 (after IGF-1-induced internalization) with reduction over time, concurrent with increased plasma membrane IGF-1R signals. In EHD1-KO cells, a more persistent co-localization is seen over time with lesser increase in plasma membrane signals over time. Top center, the data is expressed as a % of fluorescence intensity of plasma membrane IGF-1R using ImageJ. Top right, % colocalization of IGF-1R with RAB11 over time, by Pearson’s correlation coefficient quantification using ImageJ. 50 cells from three independent experiments were analyzed. **(C)** EHD1-KO impairs the Golgi to plasma membrane traffic of IGF-1R. The TC71-NTC and EHD1-KO cells pre-treated with IGF-1 (100 ng/ml) for 16 hours to deplete the cell surface IGF-1R (time 0) were subjected to serum/IGF-1 deprivation for 6,12 or 24h. Fixed and permeabilized cells were co-stained for IGF-1Rβ (green), GM130 (Golgi marker; red) and nuclei (DAPI, blue), and analyzed using confocal imaging to assess the delivery of newly-newly-synthesized IGF-1R at the Golgi followed by its delivery to the plasma membrane. Top left, a schematic of the treatments. Bottom, Co-staining for IGF-1R and GM130. The zoomed in panels (4^th^ columns for each cell line) show a small GM130-colocalizing pool of IGF-1R in TC71-NTC cells with time-dependent increase in its cell surface pool. EHD1-KO cells show an increase in the GM130-colocalizing pool of IGF-1R over time with essentially no increase in the cell surface IGF-1R. Top right, quantification of the percentage of IGF-1R fluorescence signals at the plasma membrane using ImageJ. 80 cells were analyzed from three independent experiments. B and C, scale bar, 10 μm. Mean +/− SEM. (*p<0.05, **p<0.01, ns= not significant).

To assess if EHD1 also functions as a positive regulator of the Golgi to cell surface transport of newly synthesized IGF-1R, as we reported with CSF-1R and EGFR ^4, 5^, we first treated TC71 or A673 cell lines with IGF-1 to maximally deplete the cell surface and total IGF-1R (due to ligand-induced degradation). We then switched the cells to serum/IGF-1-deprivation medium and used confocal imaging to assess the appearance of newly synthesized IGF-1R in the Golgi compartment (co-staining with the Golgi marker GM130) and at the cell surface, with quantification of the latter. At time zero (after switching to serum/IGF-1-deprivation medium), both control and EHD1-KO cells exhibited weak overall and cell surface IGF-1R signals; the cell surface IGF-1R staining progressively increased in control cells with a minor intracellular pool colocalizing with GM130 (Fig. 5C, Supplementary Fig. S7B). In contrast, only a minor increase in the cell surface pool of IGF-1R was observed over time in EHD1-KO cells; on the other hand, the KO cells exhibited strong intracellular IGF-1R persistently localizing in the GM130+ Golgi compartment (Fig. 5C, Supplementary Fig. S7B).

The marked decrease in the cell surface and total IGF-1R levels, without any change in IGF-1R mRNA levels in EHD1-depleted cells, suggested that IGF-1R is targeted for degradation. Based on our findings with CSF-1R in bone marrow-derived macrophages ^4^, we assessed if this reflected the mistargeting of IGF-1R to lysosomes upon EHD1 depletion. Treatment of steady-state cultures of Control and EHD1-KO EWS cell lines with Bafilomycin-A1, a lysosomal proton pump blocker, led to a dramatic recovery of the low total IGF-1R levels in EHD1-KO cells, nearly approaching the levels in the untreated or Baf-A1-treated control EWS cells; Baf-A1 treatment had an insignificant effect on IGF-1R levels in control cells (Fig. 6A-B). Consistent with the WB findings, confocal imaging revealed that while the pool of IGF-1R localized to LAMP1+ lysosomes in control cells was relatively unchanged upon Baf-A1 treatment, a marked and significant increase in this pool was evident in Baf-A1-treated vs. untreated EHD1-KO EWS cells (Fig. 6C-F). Collectively, these results suggest that EHD1 is required for efficient transport of IGF-1R from the Golgi and endosomal recycling compartment to the plasma membrane and that loss of EHD1 results in mistargeting of the cell-surface destined IGF-1R to the lysosome for degradation.

**Figure 6.**
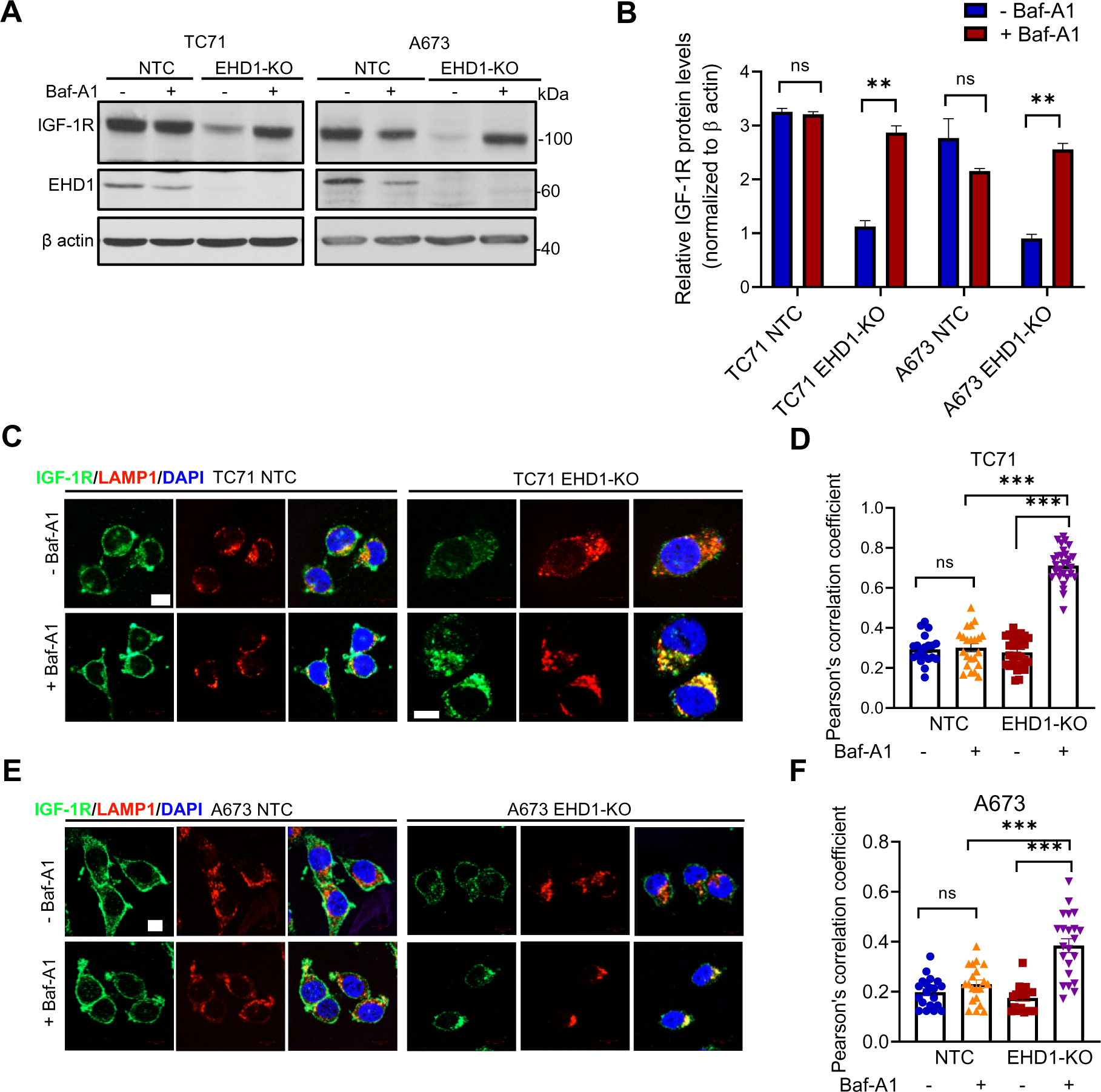
Loss of EHD1 expression leads to lysosomal degradation of IGF-1R. (A-B) Recovery of IGF-1R protein levels upon inhibition of lysosomal protein degradation with Bafilomycin A1. NTC or EHD1-KO TC71 and A673 cell lines were switched to low serum/IGF-1 free medium for 6 h in the absence or presence of Bafilomycin-A1 (200 nM) and total IGF-1R levels in cell lysates were analyzed by western blotting (A). The quantified IGF-1R signals normalized to β actin loading control are shown in B. Data represent mean +/− SEM of 3 experiments. Note the significant increase (**, p<0.01) in IGF-1R levels in EHD1-KO cells with no significant change in NTC cells (ns, not significant). **(C-F)** Lysosomal mis-targeting of IGF-1R in EHD1-KO EWS cells. NTC or EHD1-KO TC71 and A673 cells were left untreated or treated with bafilomycin-A1 as in (A-B) and co-stained for IGF-1R (green), LAMP1 (lysosome marker, red) and nuclei (DAPI, blue). IGF1-R localization to lysosomes (yellow) is visualized in merged images (third columns) in the representative images shown in C and E. Scale bar, 10 μm. Pearson’s correlation coefficients (D and F) of the co-localized IGF-1R and LAMP1 fluorescence signals were determined from analyses of n>30 cells per group from three independent experiments (**p<0.01, ***p<0.001, ns= not significant).

### IGF-1R signaling is required for EHD1 to promote the oncogenic behavior of EWS cells

Since optimal cell surface expression is essential for ligand-induced activation of RTKs ^18^, and IGF-1R activation is critical for it to promote oncogenesis and metastasis ^49^, we postulated that the positive role of EHD1 to promote the oncogenic behavior of EWS cells reflects the enhancement of IGF-1R signaling. Indeed, while control TC71 or A673 cells exhibited robust and relatively sustained IGF-1-induced phosphorylation of IGF-1R itself and of nodal readouts of its downstream signaling through AKT and MAPK signaling pathways (phospho-AKT-Ser473 and phospho-ERK1/2-Thr202/Tyr204), these responses were drastically and significantly impaired in EHD1-KO EWS cells (Fig. 7A-D). Gene-set enrichment (GSE) analysis of the RNA-seq data showed significant enrichment for genes involved in PI3K-AKT-mTOR signaling, further supporting the premise that EHD1 regulates IGF-1R signaling to promote oncogenesis (Fig. 7E). Indeed, IGF-1-dependent cell proliferation and migration were drastically and significantly reduced in EHD1-KO TC71 and A673 cell lines compared to their controls (Fig. 7F-G). Furthermore, while the IGF-1R inhibitor Linsitinib significantly reduced the IGF-1-induced proliferation and migration of control EWS cell lines, the combination of EHD1-KO and Linsitinib produced an even greater reduction in these responses (Fig. 7F-G). Flow cytometric analysis of annexin-V/PI co-stained cells revealed a significantly higher proportion of apoptotic cells in EHD1-KO EWS cell lines; Linsitinib significantly increased the proportion of early and late apoptotic cells in control EWS cells and more so in EHD1-KO TC71 and A673 cells (Fig. 7H, Supplementary Fig. S8A). The additional Linsitinib inhibition of IGF-1-induced oncogenic traits in EHD1-KO cell lines is consistent with lower residual levels of IGF-1R in these cells.

**Figure 7.**
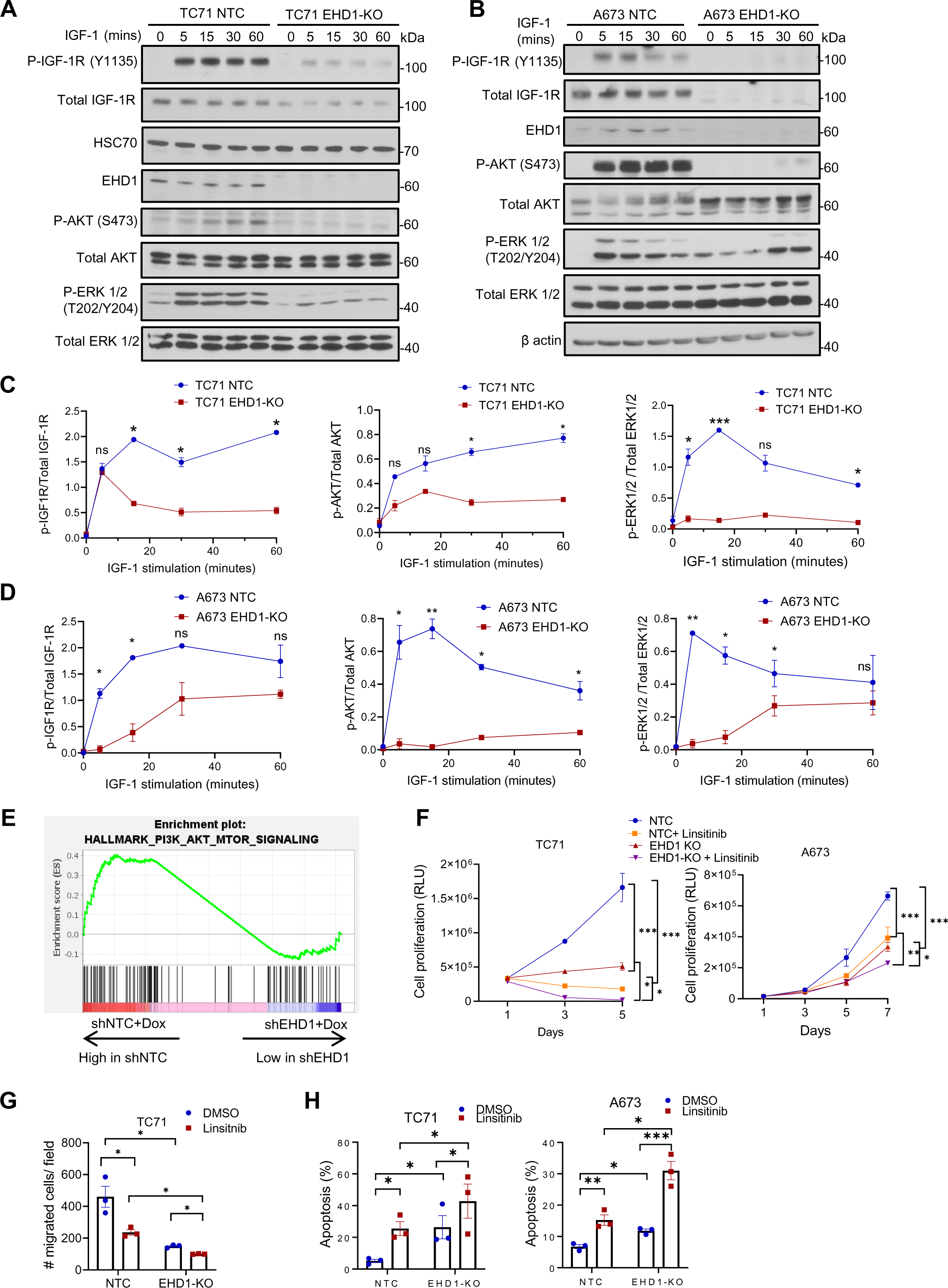
Loss of EHD1 expression in EWS cells impairs the IGF-1-dependent signaling downstream of IGF-1R. (A-B) Western blot analysis of phosphorylation of IGF-1R and key signaling pathway reporters (AKT and ERK1/2). NTC or EHD1-KO TC71 and A673 cell lines were pre-starved for 24h in low serum/IGF-free medium and left unstimulated (0) or stimulated with IGF-1 (50 ng/ml) for the indicated time points (min, minutes). Cell lysates were analyzed by Western blotting with the indicated antibodies, with β actin as loading control. **(C-D)** Densitometric quantification of the phosphorylation signals of IGF-1R, AKT and ERK (from the data represented in A-B) normalized to the values of total proteins. Data represent mean +/− SEM of 3 experiments. **(E)** Gene-set enrichment (GSE) analysis from RNA-sequencing of two groups of TC71 cell lines-TC71 shEHD1+Dox vs. shNTC+Dox, showing enrichment of PI3K-AKT-mTOR signaling genes in shNTC+Dox cells and significant downregulation of the same in the shEHD1+Dox group. **(F)** EHD1-KO impairs the IGF-1-dependent pro-survival effects in EWS cells. Flow cytometric analysis of apoptosis in the indicated cells treated with or without 1 μM linsitinib for 24 hours as assessed by Annexin-V and PI staining. **(G)** Impaired IGF-1-induced proliferation in EHD1-KO EWS cell lines. NTC or EHD1-KO TC71 and A673 cells were cultured in regular medium for 24h, switched to medium with 1% FBS and 100 ng/ml IGF-1) in the absence or presence of 1 μM IGF-1R inhibitor linsitinib and cell proliferation measured at the indicated time points by Cell-Titer Glo assay. Data represent mean +/− SEM of three experiments, each in six replicates. **(H)** Impaired IGF-1-induced cell migration in EHD1-KO EWS cell lines. NTC or EHD1-KO TC71 and A673 cells plated in top chambers of trans-wells in the absence or presence of 1 μM IGF-1R inhibitor linsitinib, with migration towards the medium with 1% FBS and 100 ng/ml IGF-1 in lower chambers. Data represent mean +/− SEM of three experiments; *, p<0.05, **p<0.01***, p<0.001.

To directly assess the requirement of IGF-1R for EHD1-dependent elevation of the oncogenic behavior of EWS cells, we targeted IGF-1R by multiple approaches in mEHD1-overexpressing SK-ES-1 cell line, which exhibits a specific EHD1 overexpression-dependent enhancement of oncogenic traits (Fig 8A-B, Supplementary Fig S8E). While control siRNA transfection had no impact on IGF-1-induced cell proliferation, migration, or invasion, siRNA KD of IGF-1R, pharmacological inhibition with Linsitinib, or treatment with an inhibitory monoclonal antibody 1H7 ^50, 51^ impaired these *in vitro* readouts to a level comparable to those in parental EHD1-low cells (Fig. 8C-I, Supplementary Fig S8B-G). Comparable results were observed when apoptosis was measured as a readout (Fig. 8D, Supplementary Fig S8B).

**Figure 8.**
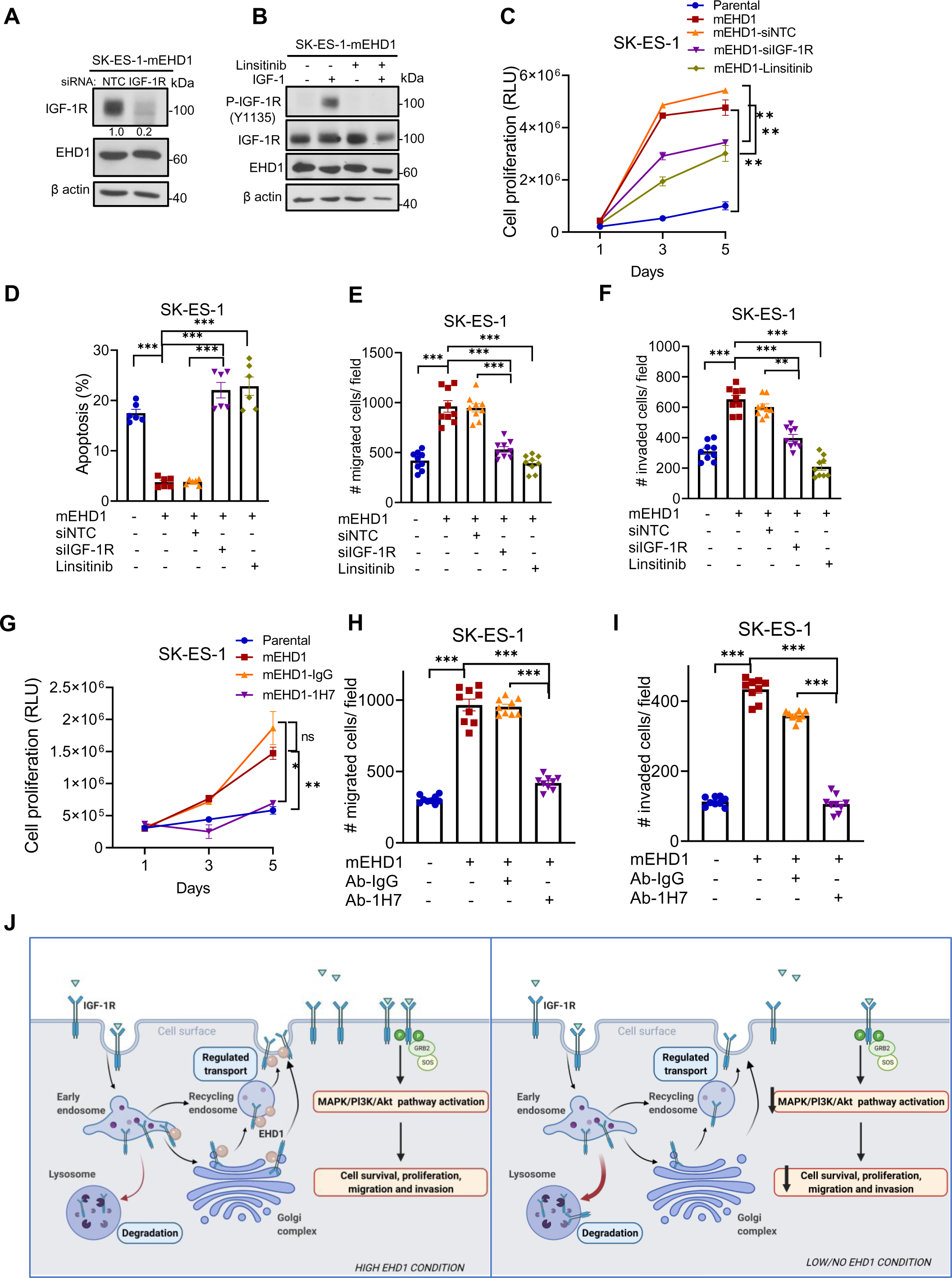
EHD1-dependent upregulation of oncogenic attributes of EWS cell lines requires the IGF-1R. Mouse EHD1 (mEHD1)-overexpressing SK-ES-1 cell line was used to assess the requirement of IGF-1R in EHD1-driven pro-oncogenic attributes. SK-ES-1-mEHD1 cells transiently transfected with non-targeting control (NTC) or IGF-1R-targeted siRNA or treated with IGF-1R inhibitor linsitinib (1 µM), IGF-1R mAb 1H7 (5 µg/mL), mouse Isotype IgG1 (5 µg/mL) were studied for indicated traits. **(A)** Representative western blot confirming the effective IGF-1R knockdown upon transient IGF-1R siRNA relative to NTC siRNA transfection. **(B)** Western blot showing effective elimination of phospho-IGF-1R signals by IGF-1R siRNA knockdown or linsitinib treatment in SK-ES-1-mEHD1 cells. **(C)** Elevated cell proliferation upon mEHD1 overexpression requires IGF-1R expression and activity. The SK-ES-1-mEHD1 cells were analyzed for IGF-1 (100 ng/ml)-dependent cell proliferation by Cell-Titer Glo assay with or without the indicated treatments. Parental SK-ES-1 cells without any treatments provided a baseline of cell proliferation without mEHD1 overexpression. **(D)** Elevated cell survival upon mEHD1 overexpression requires IGF-1R expression and activity. The SK-ES-1-mEHD1 cells grown in the presence of IGF-1 (100 ng/ml) without or with the indicated treatments for 3 days were analyzed for the proportion of apoptotic cells by FACS after Annexin-V and PI staining. **(E-F)** Elevated cell migration and invasion upon mEHD1 overexpression requires IGF-1R expression and activity. The SK-ES-1-mEHD1 cells were analyzed for IGF-1 (100 ng/ml)-dependent trans-well cell migration or invasion without or with the indicated treatments. Parental SK-ES-1 cells without any treatments provided a baseline of cell migration without mEHD1 overexpression. Mean +/− SEM of 3 experiments, each in triplicates. *p<0.05; **p<0.01; ***p<0.001; ns, not significant. **(G-I)** Inhibition of cell proliferation **(G),** cell migration**(H)** and invasion**(I)** in SK-ES-1 mEHD1 cells with IGF-1R mAb 1H7. Analyses done as in C-F. **(J)** A model of how the EHD1/IGF-1R axis promotes the IGF-1R-mediated signaling and tumor progression in Ewing Sarcoma. EHD1 overexpression enhances the endocytic recycling and Golgi to plasma membrane transport of IGF-1R to elevate the cell surface receptor levels, thus enhancing IGF-1R-dependent signaling. Loss of EHD1 leads to IGF-1R mistargeting to lysosomes where it is degraded, resulting in reduced cell surface IGF-1R, diminished IGF-1R signaling and impaired tumorigenesis.

## 4. DISCUSSION

Besides driver oncogenes, tumor cells turn on multiple adaptive pathways for successful primary tumor growth and metastasis. Delineating these oncogenesis-enabling pathways is likely to identify novel biomarkers of malignant behavior and therapeutic responses of tumors and in some cases, offer opportunities for therapeutic targeting. Here, using Ewing Sarcoma (EWS) as a tumor model, we demonstrate that the intracellular vesicular traffic regulatory protein EHD1 promotes tumorigenesis and metastasis by serving as a required element of IGF-1R traffic to enable IGF-1R-mediated oncogenic programs. While EWS is a relatively uncommon malignancy, it is the second most common bone and soft tissue tumor of children and young adults ^22^. Importantly, the novel mechanistic insights we uncover using EWS models are likely to be broadly relevant to malignancies where RTKs serve as drivers or enablers of oncogenesis, and EHD1 protein is overexpressed.

In a large EWS tumor panel, we found moderate to high EHD1 overexpression in nearly 90% of patients, with significantly higher levels in metastatic tumors (Fig.1D-E). Query of publicly-available data revealed the high EHD1 mRNA expression to be associated with shorter patient survival (Fig.1A-B). Thus, clinical data support a positive role of EHD1 protein in EWS tumorigenesis. These findings are consistent with reports of EHD1 overexpression in other cancers, in many cases associated with shorter patient survival or resistance to therapy ^8, 11, 16^.

Our comprehensive genetic analyses of EWS cell models definitively demonstrate that EHD1 propels tumorigenic and metastatic behavior in EWS. Use of Doxycycline-inducible shRNA knockdown in cell line models demonstrated a strong dependence of cell proliferation, tumorsphere growth, cell migration, and invasion on EHD1 (Fig.2C-H), with a stronger impact in cells lines with higher EHD1 expression (A673 and TC71) and a more modest impact in cells (SK-ES-1) with lower EHD1 levels (Supplementary Fig.S1B, G). Reciprocally, ectopic mouse *Ehd1* overexpression in the latter cells markedly enhanced their pro-tumorigenic and pro-metastatic traits (Fig.2J-M). EHD1-KO in A673 and TC71 cell models confirmed the requirement of EHD1 for the *in vitro* pro-tumorigenic and pro-metastatic behavior of EWS cells, and re-expression of mEHD1 restored the EHD1-KO defects (Fig.2I). Use of Dox-inducible KD or EHD2-KO EWS cell models in a bone implant model in nude mice demonstrated a key role of EHD1 in EWS tumorigenesis and metastasis *in vivo,* and the defective tumorigenic ability of EHD1-KO cells was completely restored by mEHD1 rescue (Fig.3A-B). Furthermore, the modest metastases forming ability of parental EWS cells was completely abolished by EHD1-KO; notably, the mEHD1-rescued EHD1-KO cells, which express higher EHD1 levels than the parental cells, showed significantly more metastatic growths (Fig.3C-D). A hallmark of bone-associated tumors is the destruction of the surrounding bone ^52^. Indeed, compared to significant bone destruction by parental cell implants, EHD1-KO cells failed to do so, and the process was accentuated in mEHD1-rescued KO cells (Fig.3E-F). Collectively, our clinical-pathological studies combined with our *in vitro* and *in vivo* genetic perturbation studies provide compelling evidence for a key role of EHD1 overexpression in sustaining EWS tumorigenesis and metastasis.

Our studies provide novel insights into how EHD1 serves in a pro-tumorigenic and pro-metastatic role. Our mechanistic studies were focused on two key considerations, one the established role of EHD1 in regulating intracellular traffic of multiple cell surface receptors ^1, 53, 54^, and our previous studies that have established a key role of EHD1 to ensure high cell surface expression of RTKs by regulating key aspects of their traffic ^4, 5^. Our unbiased query of human receptor tyrosine kinome identified IGF-1R as a specific target (Fig.4A). Our comprehensive cell biological analyses demonstrate that EHD1 is required for Golgi to plasma membrane traffic of newly-synthesized IGF-1R to ensure high pre-activation levels of total and cell surface IGF-1R (Fig.5C), the latter a requirement for subsequent ligand-induced activation of signaling and cellular responses ^32^. In addition, EHD1 plays a positive role in post-activation recycling of IGF-1R to help return it to the cell surface (Fig.5B), uncovering a second trafficking mechanism known to help sustain cell surface RTK levels by countering their lysosomal targeting ^20^. Consistent with the key roles of EHD1 in regulating IGF-1R traffic to sustain its cell surface expression while negating its lysosomal degradation, our biochemical and subcellular localization analyses establish that lack of EHD1 leads to marked mistargeting of IGF-1R to lysosomes where it is degraded (Fig. 6). Previous analyses have shown that EHD1 can interact with IGF-1R ^48^, which we find is also the case in EWS cell models (Fig. 5A), but a role for EHD1 in regulating IGF-1R traffic has not been shown previously.

Notably, ligand-induced internalization, lysosomal degradation, and recycling of IGF-1R are well-established aspects of its traffic and signaling ^55–58^. Post-endocytic recycling of IGF-1R has been shown to be positively regulated by myoferlin ^59^, RAB11-FIP3 ^60^, and GIGYF1 ^61^. Thus, our studies identify EHD1 as a new regulator of IGF-1R endocytic recycling. RAB11-FIP3 is a component of endocytic recycling, in which EHD1 plays a key role ^1^, and a family member RAB11-FIP2 interacts with EHD proteins ^47^, suggesting the possibility that EHD1 may function together with RAB11-FIP proteins to regulate the recycling of IGF-1R and potentially other RTKs. Interestingly, ligand-dependent IGF-1R localization to Golgi has been associated with the migratory behavior of tumor cells, suggesting signaling capabilities of the Golgi-localized receptor ^62^. In previous studies, we found EHD1 to play a role in retrograde traffic of cell surface EGFR to Golgi ^5^, suggesting the possibility that EHD1 could play a similar role in IGF-1R traffic.

In contrast to its post-activation traffic, mechanisms that regulate the availability of IGF-1R at the cell surface prior to ligand binding have been less explored. Interestingly, Smoothened was found to positively regulate IGF-1R levels in lymphoma and breast cancer cell lines by stabilizing it in plasma membrane lipid rafts and preventing its lysosomal targeting ^63^. Whether Smoothened regulates endocytic recycling or Golgi to cell surface IGF-1R traffic was not explored. Notably, we have shown EHD1 regulation of Smoothened traffic in primary cilia ^64^, raising the possibility that Smoothened and EHD1 may co-regulate IGF-1R traffic.

Our findings linking EHD1 overexpression to regulation of an RTK well-established to control multiple aspects of oncogenesis provided a plausible basis for EHD1’s pro-oncogenic role we uncovered. We provide multiple lines of evidence that this indeed is the case. Reduced cell surface IGF-1R expression upon EHD1-KO directly translated into reduced activation of downstream signaling (Fig.7A-D), and transcriptomic analyses support this conclusion (Fig.7E). Accordingly, EHD1-depleted EWS cells showed markedly reduced IGF-1-dependent proliferation, survival, and migration (Fig.7F-H). Furthermore, EHD1-KO status sensitized the EWS cells to elevated levels of apoptosis and a further reduction in cell migration upon inhibition of IGF-1R with Linsitinib (Fig.7F-H). Thus, our analyses clearly establish that EHD1 overexpression, by sustaining elevated levels of total and cell surface IGF-1R, promotes multiple aspects of oncogenesis in EWS. While signaling through IGF-1R is well established to promote oncogenesis in EWS ^35^, we directly establish that elevation of IGF-1R levels and subsequent IGF-1R-mediated signaling underlies the ability of EHD1 to promote the oncogenic behavior of EWS cells. Using the mEHD1-overexpressing SK-ES-1 cell model of EHD1-driven elevation of oncogenic behavior (Fig. 1J-M), our multi-pronged studies using siRNA KD, kinase inhibition, and an inhibitory antibody approach demonstrates a requirement of IGF-1R for EHD1 overexpression-driven oncogenic traits (Fig.8A-I). Thus, our studies clearly establish the upregulation of IGF-1R levels and signaling by overexpressed EHD1 as a key oncogenic adaptation in EWS. Consistent with this conclusion, analysis of a large cohort of EWS patient samples showed a significant positive correlation between EHD1 and IGF-1R protein levels (Fig. 8F-H).

Our findings using an EWS model have potential implications for the pro-oncogenic role of EHD1 and RTK-dependent sustenance of tumorigenesis and metastasis in other cancers. EHD1 overexpression is linked to shorter survival and chemotherapy/EGFR-TKI resistance in NSCLC ^16^, apparently through PI3K-AKT pathway activation by interaction with the microtubule protein TUBB3 and stabilization of microtubules ^16^ and through promotion of aerobic glycolysis via a 14-3-3z-dependent b-catenin-c-Myc activation pathway ^14^. While these mechanisms may operate independently of RTK signaling, the key roles of the wildtype or mutant EGFR, as well as IGF-1R and other RTKs, in NSCLC pathogenesis and therapeutic resistance ^65^ raise the possibility that EHD1 overexpression activates these pathways by sustaining RTKs, as we show in the EWS model. Association of EHD1 overexpression with EGFR-TKI resistance in NSCLC ^11, 16^ and with higher expression of EGFR, phospho-EGFR and RAB11-FIP3 ^12^ support this idea.

While our studies focus on the linkage of EHD1 with an RTK, EHD1 overexpression may also regulate other oncogenesis-related cell surface receptors, given its broader roles. Indeed, EHD1 overexpression was shown to promote cancer stem cell-like traits in glioblastoma and lung cancer by promoting CD44 recycling while suppressing its degradation ^15, 66^, promote cisplatin resistance in NSCLC by regulating cisplatin accumulation in cells, presumably by regulating transporter levels ^9^, and potentiate angiogenesis by promoting b2 adrenergic receptor recycling ^13^. Cell biological studies have also shown a positive role of EHD1 in β1 integrin recycling ^53^. Future studies of the kind described here in the context of an RTK, IGF-1R, should help uncover the individual or combined roles of the various EHD1-regulated cell surface receptors in promoting tumorigenesis and metastasis.

In conclusion, our analyses in an EWS tumor model show that EHD1 overexpression promotes oncogenesis by post-translationally upregulating the trafficking itinerary of an RTK, IGF-1R.

## AUTHOR CONTRIBUTIONS

Designing research studies: S.C., H.B., V.B., B.C.M. Conducting experiments: S.C., A.M.B., I.M., H.L., B.C.M. Acquiring and analyzing data: S.C., B.C.M., A.K., S.M., N.C., J.A.L., A.L.B., I.M., J.L.M, D.W.C., J.M.R. Providing reagents: M.D.S., G.G. Writing the manuscript: S.C., H.B. All authors have read and agreed to the version of manuscript.

## ACKNOWLEDGEMENTS

This research was funded by: Pilot grants from the Pediatric Cancer Research Program of the Children’s Hospital Research Institute, UNMC; Pilot grants from the Fred & Pamela Buffett Cancer Center (HB & VB); Department of Defense grants W81XWH-17-1-0616 and W81XWH-20-1-0058 to HB and W81XWH-20-1-0546 to VB; the NIH grants R21CA241055 and R03CA253193 to VB; the Raphael Bonita Memorial Fund; and support to UNMC core facilities from the NCI Cancer Center Support Grant (P30CA036727) awarded to Fred & Pamela Buffett Cancer Center and from the Nebraska Research Initiative. SC and AMB received the University of Nebraska Medical Center Graduate Student Fellowships.

---

**Supplementary Fig. S1.**
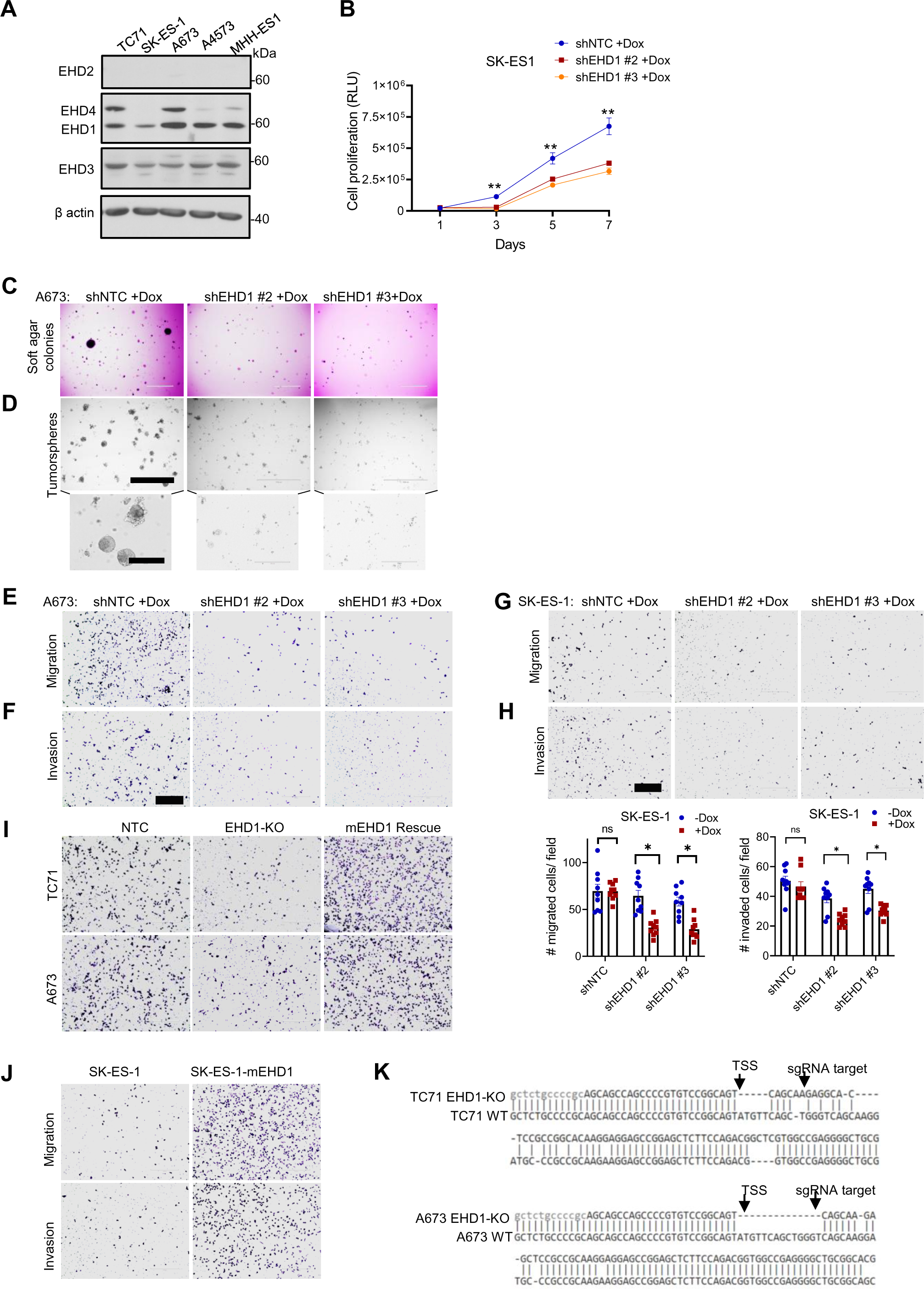
EHD1 is required to sustain the in vitro oncogenic traits of Ewing Sarcoma cell lines (A) Immunoblot analysis of EHD1 and its family members in the indicated EWS cell lines. β actin served as loading control. **(B**) Impaired proliferation of SK-ES-1 cells upon EHD1 knockdown. Cell-titer-glo assay at the indicated time points in the presence of Dox. Mean +/− SEM of three experiments, each with six replicates. **(C-D)** Impaired soft agar colony growth (C) and tumor-sphere forming ability (D) of A673 cells upon EHD1 knockdown. Representative images of A673 cells (Fig. 2E-F; bottom) are shown. **(E-F)** Impaired trans-well migration (E) and invasion (F) in A673 cells upon EHD1 knockdown. Representative images of A673 cells (Fig. 2G-H; bottom) are shown. **(G-H)** Impaired of trans-well migration (G) and invasion (H) in SK-ES-1 cells upon EHD1 knockdown. Top, representative images of SK-ES-1 cells; scale bar, 400 μm. Bottom, quantification of the number of migrated/invaded cells per high-power field; mean +/− SEM of three experiments each in triplicates. **(I)** Representative images of Fig2I showing impaired trans-well cell migration upon EHD1 knockout (KO) and rescue of migration defect by mEHD1. **(J)** Representative images of Fig2L-M showing increase in transwell migration and invasion upon mEHD1 overexpression in SK-ES-1 cells. **(K)** CRISPR-Cas9 knockout site assessment of EHD1 by Sanger sequencing showing deletion of bases, removal of start codon and frameshift mutations near the sgRNA targeted sequence in TC71 and A673 cell lines. Wildtype (WT) sequences are shown as reference.

**Supplementary Fig. S2.**
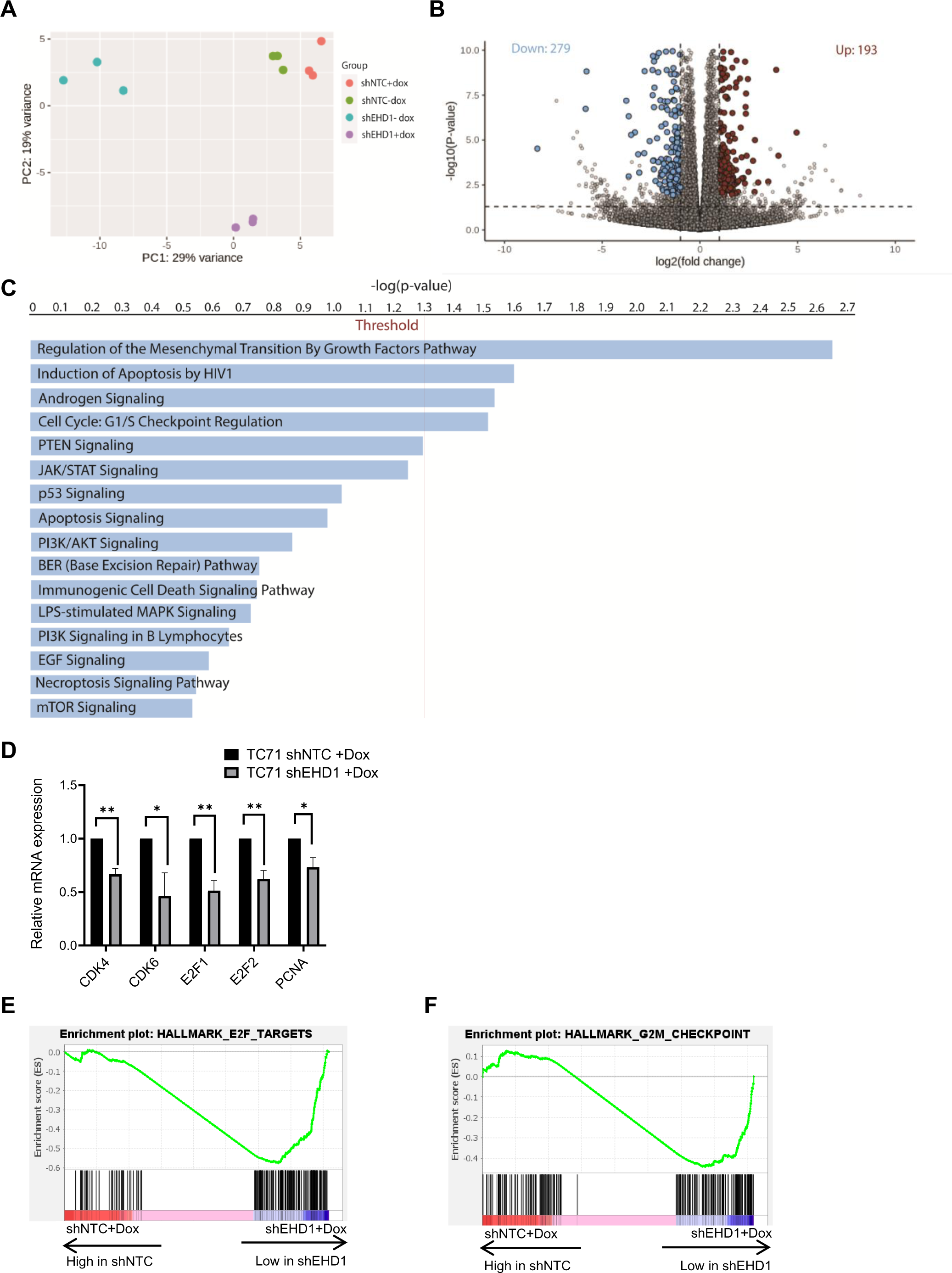
RNA-Sequencing analysis of shNTC and shEHD1+dox TC71 cell line (A) PCA analysis of RNA-seq data shows four datasets – TC71 shNTC -/+Dox, shEHD1 -/+ Dox. PC1 represent 29% variance and PC2 represent 19% variance. **(B)** Volcano plot showing differentially expressed genes – upregulated (in red), downregulated (in blue). **(C)** Canonical signaling pathways affected by the differentially expressed genes by Ingenuity-Pathway Analysis (IPA) software. Vertical line indicates threshold of -log_10_(p-value) =1.3 **(D)** Validation of G1 to S cell cycle regulatory genes by qPCR analysis in TC71 shEHD1 -/+Dox groups. (mean +/− SEM of three experiments, *p<0.05, **p<0.01) **(E-F)** Gene-set enrichment (GSE) analysis was performed on the RNA-sequencing of two groups of TC71 cell lines-TC71 shEHD1+Dox vs. shNTC+Dox, showing enrichment of E2F-targets (E) and G2-M cell cycle checkpoint(F) genes in shNTC+Dox cells and significant downregulation of the same in the shEHD1+Dox group. Differential expression was assessed by DESeq2 and significantly changed genes were required to have a Benjamini–Hochberg adjusted p-value of <0.05 and a 2-fold change in expression.

**Supplementary Fig. S3.**
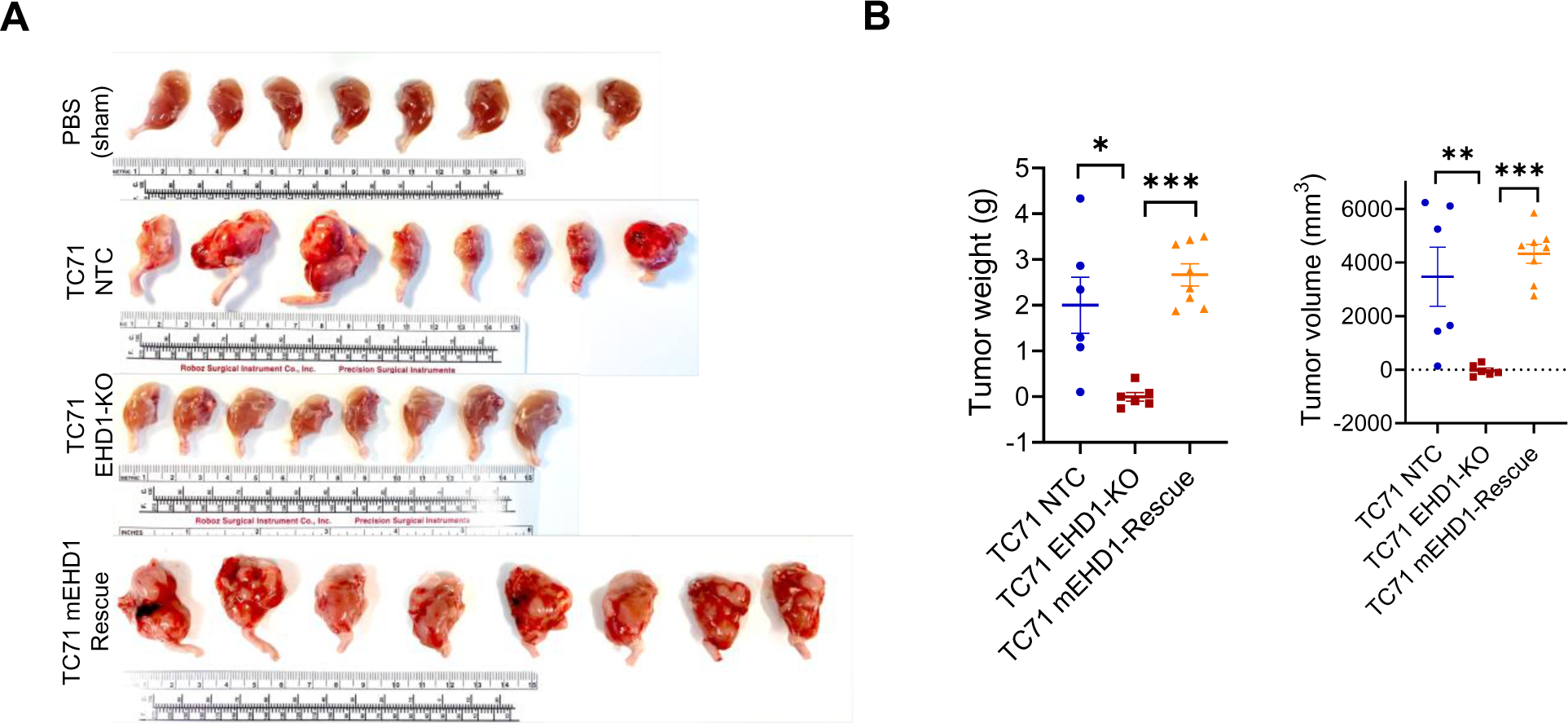
Loss of EHD1 expression markedly impairs the growth of bone implanted EWS cells. (A) Images of tumors harvested at the end of the experiment shown in Fig. 3A-D together with the sham (PBS)-injected contralateral legs of the mice injected with TC71-NTC cells. **(B)** Quantification of harvested tumor weight (left panel) and volume (measurement with calipers (volume = length x width x depth/2); right panel. Values of sham-injected legs were subtracted from the experimental values.

**Supplementary Fig. S4.**
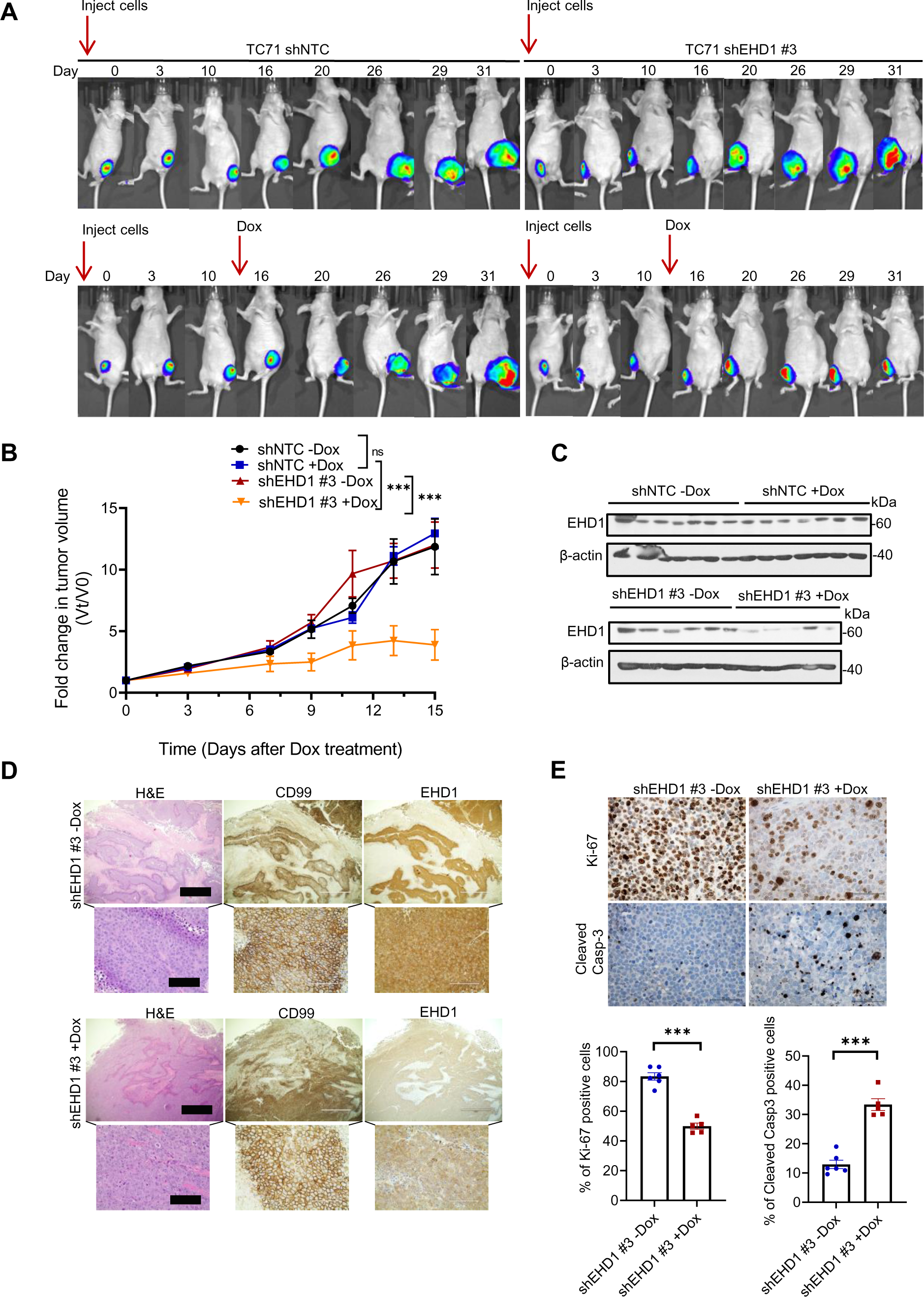
Demonstration of EHD1 requirement for EWS tumorigenesis using Dox-inducible shRNA knockdown. Intratibial tumor injections with the indicated TC71 cell lines with Dox-inducible control (shNTC) or EHD1 (shEHD1#3) shRNA were done as in Fig. 3.; 7 mice/group. **(A)** Images of one out of seven mice in various groups with super-imposed luminescence signals over 31 days. **(B)** Tumor growth with the injections of the indicated TC71 derivatives, with or without Dox administration. Differences between the indicated groups analyzed using the two-way ANOVA; ***p<0.001. Note lack of impact of Dox on tumors generated with TC71 shNTC. **(C)** Western blots of harvested tumor tissue to confirm Dox-induced EHD1 knockdown in TC71 shEHD1 #3 xenografts. **(D)** Representative tumor sections of TC71-shEHD1-Dox and shEHD1+Dox tumors stained with H&E (left panels), CD99 (middle panels, demarcating the human EWS tumor cell area) and EHD1 (right panels). **(E)** IHC staining for Ki-67 and Cleaved-Caspase-3 in tumor sections from the indicated groups. Top, representative images; bottom, quantification IHC staining positive cells. Mean +/− SEM; *p<0.05, **p<0.01, ***p<0.001, ns= not significant.

**Supplementary Fig. S5.**
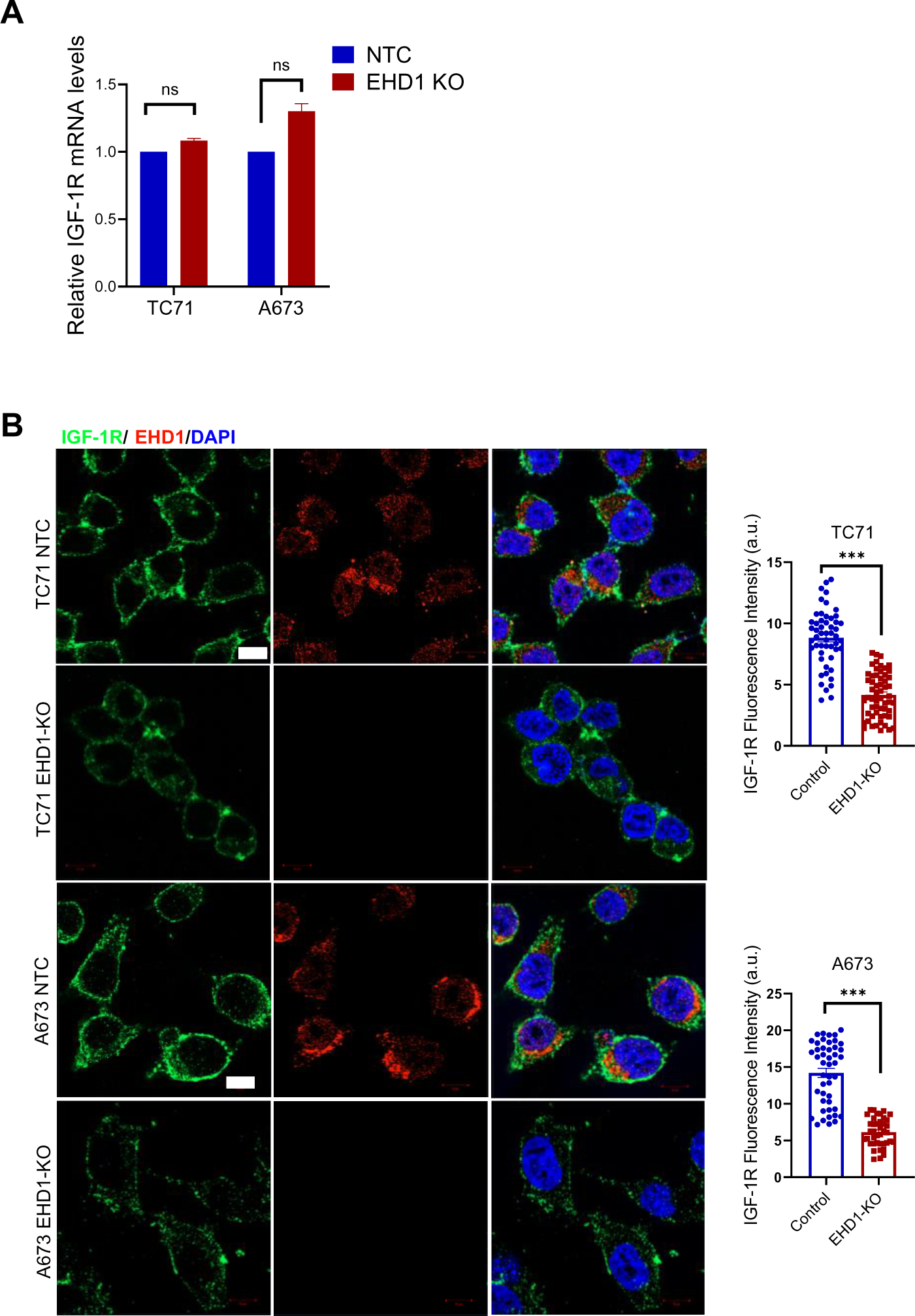
Identification of insulin like growth factor-1 receptor (IGF-1R) as a regulatory target of EHD1 in EWS. (A) EHD1-KO in EWS cell lines does not affect the IGF-1R mRNA expression. Shown are the qRT-PCR based results of IGF-1R mRNA expression, normalized to GAPDH and expressed as a fold change relative to the respective NTC control cell lines (set to 1). Data represent mean +/− SEM of 3 independent experiments (ns= not significant). (B) Reduction in IGF-1R levels upon EHD1-KO in EWS cell lines analyzed by immunofluorescence staining and confocal imaging. IGF-1R (green) and EHD1 (red) staining in Control and EHD1-KO TC71 and A673 cells. Cells grown under steady-state were fixed and permeabilized and stained with the indicated antibodies (with concurrent IgG controls; not shown). Left, representative confocal images. Merged pictures with DAPI (blue) are shown in right panels. Right, Quantification of the IGF-1R fluorescence intensity. Scale bar (only shown in left panels of the NTC lines), 10 μm. Data points represent images of 60 cells pooled from three independent experiments.

**Supplementary Fig. S6.**
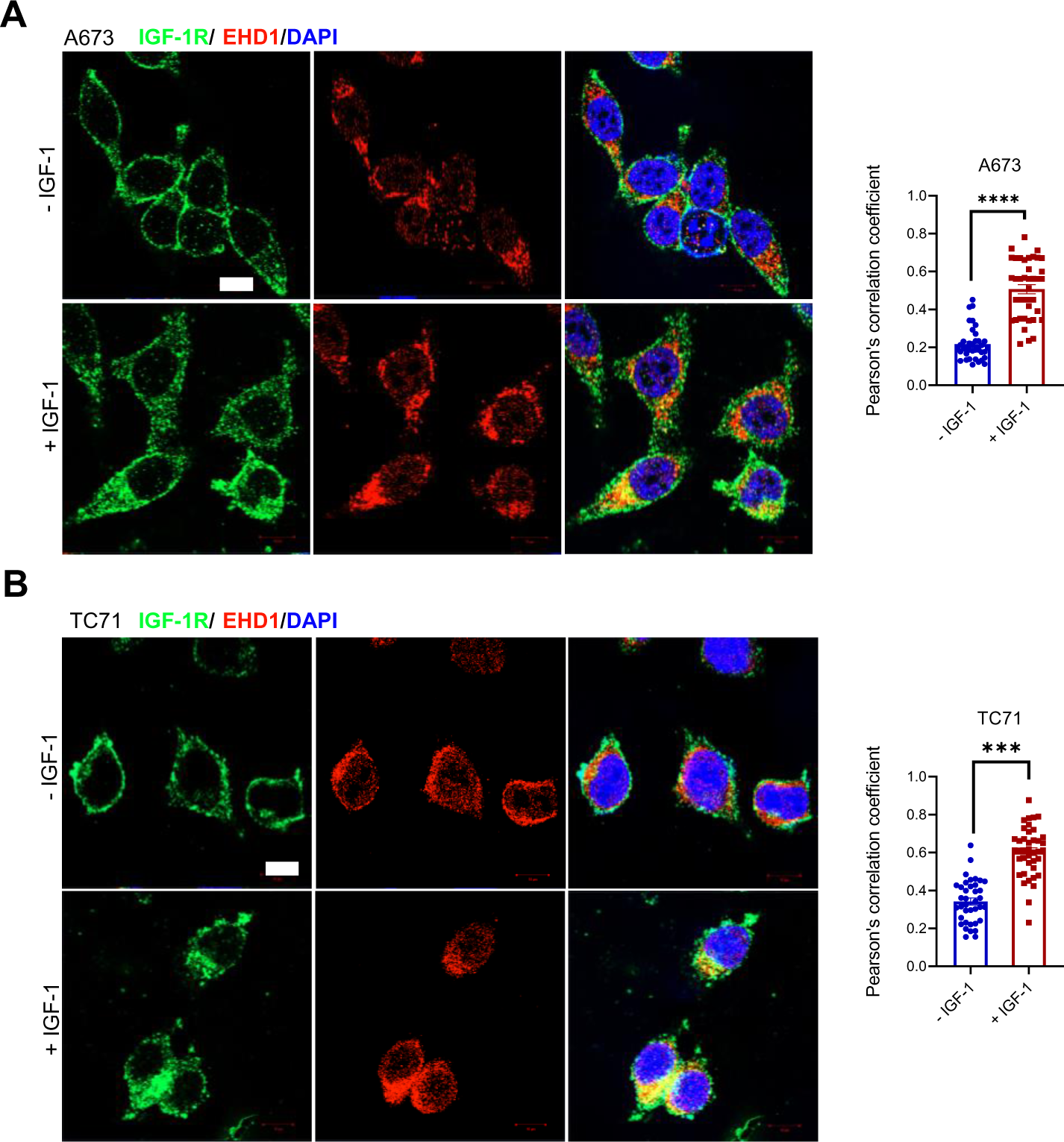
EHD1 and IGF-1R colocalize in intracellular vesicular structures. (A-B) Representative confocal images of the colocalization of EHD1 (red) and IGF-1R (green) in A673 (A) and TC71 (B) cells without (top panels) and with (bottom panels) IGF-1 (50 ng/ml) stimulation for 1h. Merged pictures (right panels) with DAPI (blue) show colocalization within perinuclear vesicular structures. Scale bar, 10 μm. Colocalization was assessed in 40 cells in three independent experiments to determine the colocalization coefficients. Data represent the mean +/− SEM. ***p<0.001.

**Supplementary Fig. S7.**
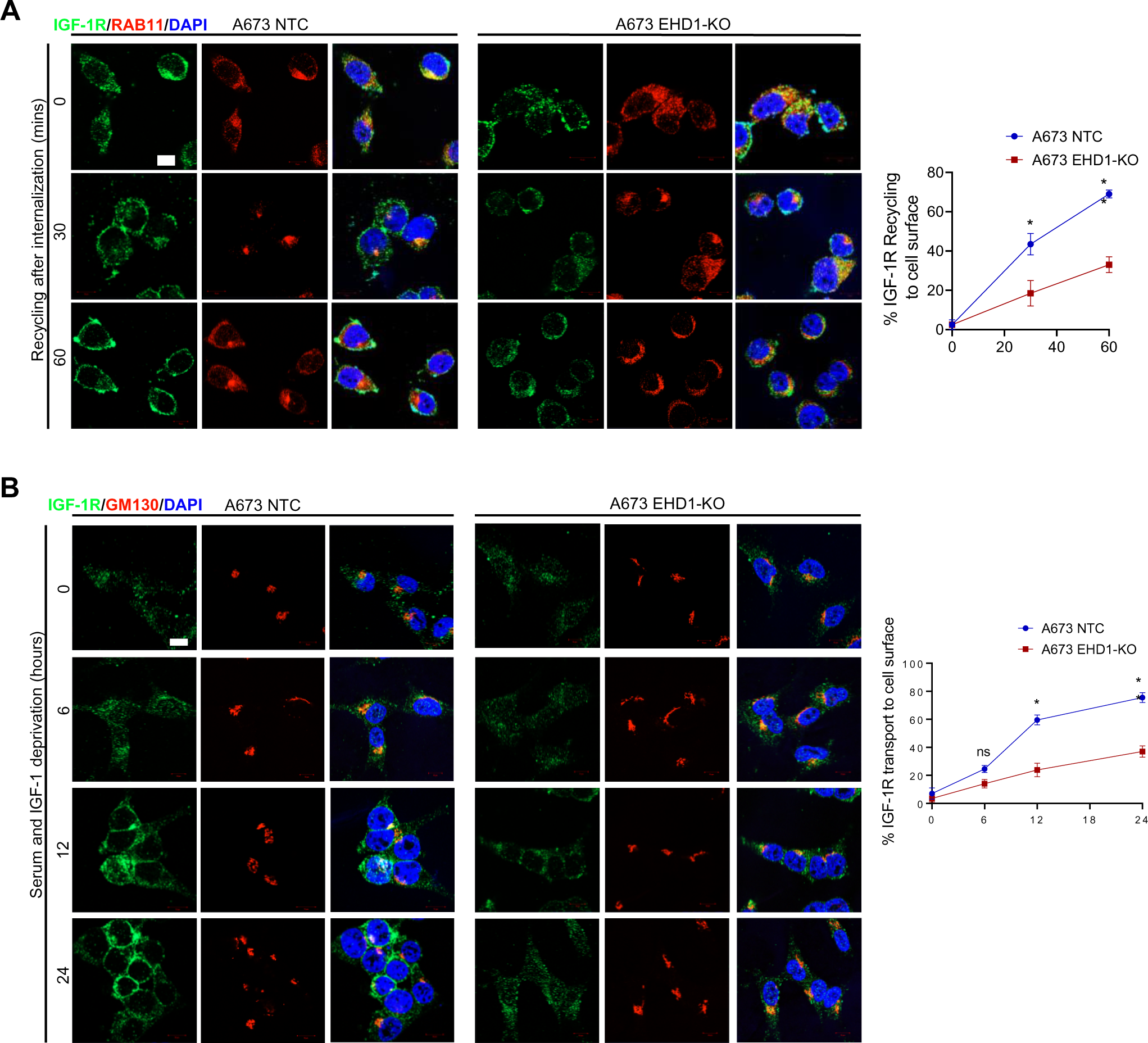
Impairment of IGF-1R transport from the Golgi and recycling endosomes to the plasma membrane by EHD1-KO in A673 cell line. The analyses with A673 NTC and EHD1-KO cells were carried out exactly as described in Fig. 5 for TC71 cells. (A) Analysis of IGF-1R endocytic recycling; IGF-1R, green; Recycling endosome (Rab11+), red. (B) Analysis of IGF-1R Golgi to plasma membrane transport; IGF-1R, green; Golgi (GM130+), red. Top panels, representative confocal images (zoomed images in third column). Bottom, quantification of cell surface IGF-1R at various time points using the ImageJ. Data represent mean +/− SEM. *p<0.05; **p<0.01; ns, not significant. Scale bar, 10 μm.

**Supplementary Fig. S8.**
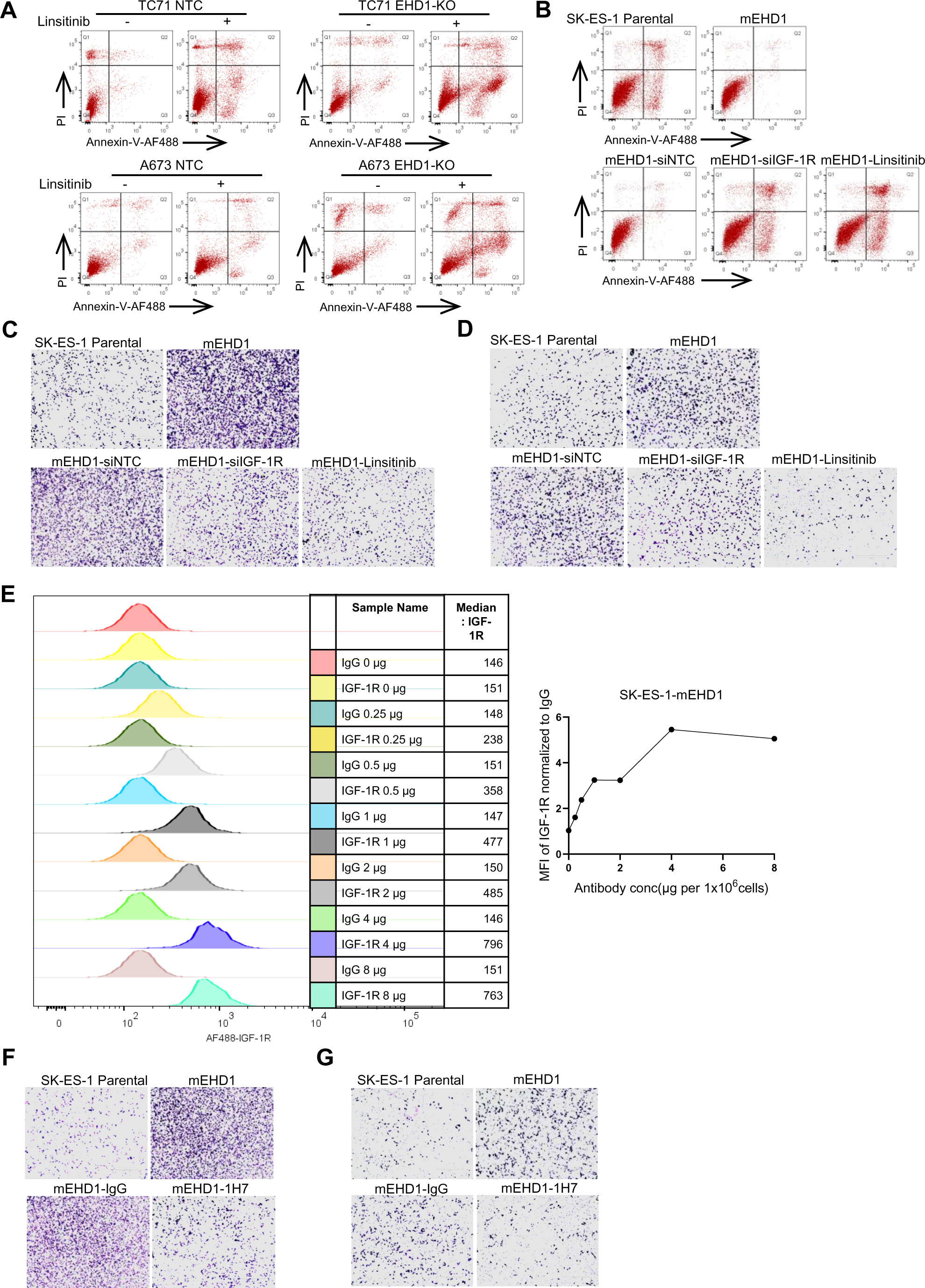
EHD1-dependent upregulation of oncogenic attributes of EWS cell lines requires the IGF-1R. (A) Representative flow panel of Annexin-V-PI assay (Figure 7F) in TC71 and A673 NTC and EHD-KO cells, with the indicated treatments. **(B)** Representative flow panel of Annexin-V-PI assay (Figure 8D) in SK-ES-1 mEHD1 overexpressing cell line, with the indicated treatments. **(C-D)** Representative high-power fields of migration and invasion assays corresponding to Figure 8E-F. **(E)** Dose-response of IGF-1R monoclonal antibody 1H7 showing saturation at 4µg antibody concentration/million cells. Representative flow panels(left), graph plotting Median fluorescence intensity (MFI) normalized to same concentration of mouse isotype control IgG1(right) **(F-G)** Representative high-power fields of migration and invasion assays corresponding to Figure 8H-I.

**Table 1:**
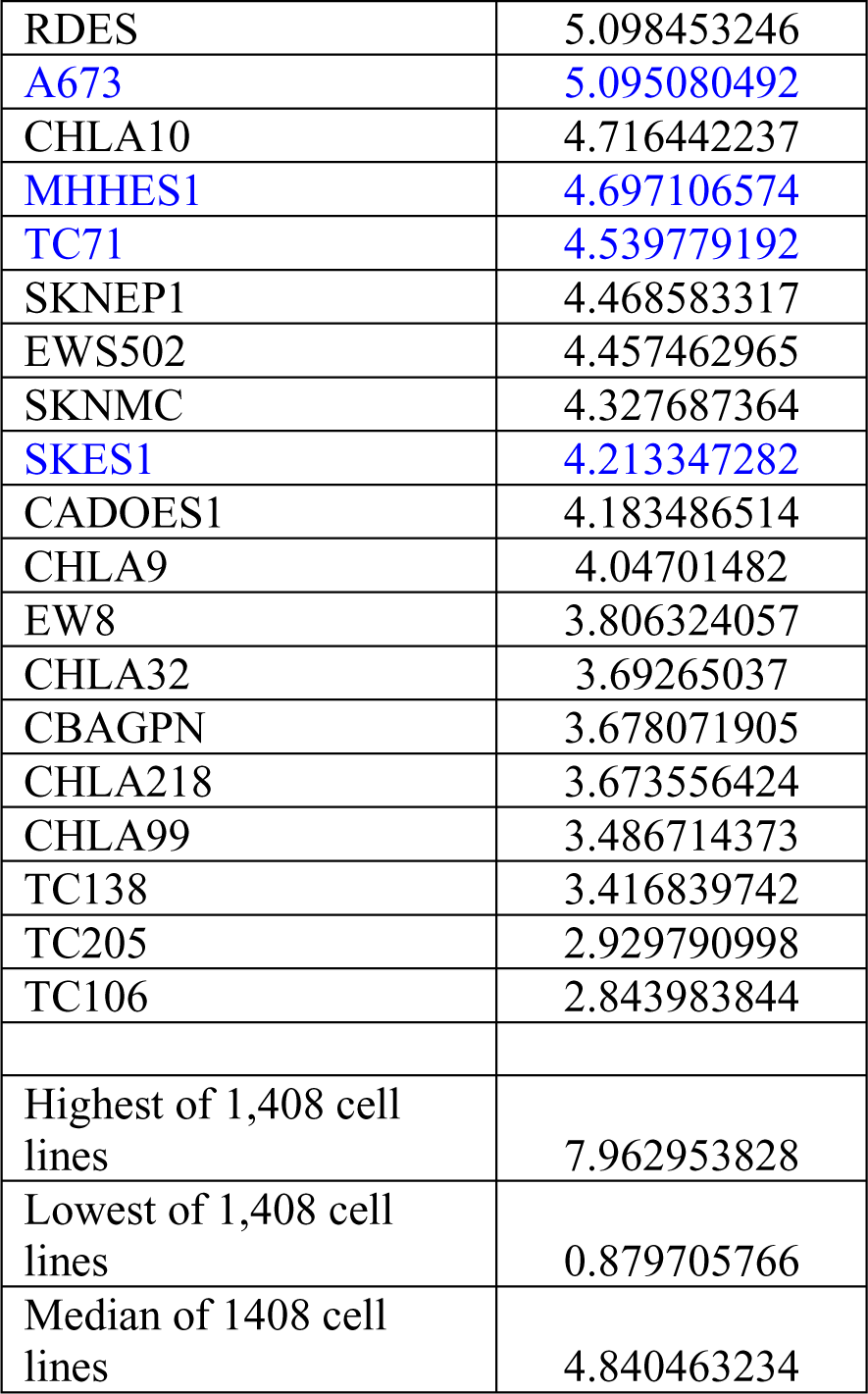
mRNA expression of EHD1 in Ewing Sarcoma cell lines (CCLE):

**Table 2.** Higher expression of EHD1 in metastatic lesions:

| EHD1 | Tissue sample |  |  |  | Total |
| --- | --- | --- | --- | --- | --- |
|  | Localized | Disseminated | Relapse | Metastasis |  |
| 0 | 33 (12.4%) | 0 (0%) | 2 (20%) | 0 (0%) | 35 |
| 1 | 118 (44.2%) | 15 (33.3%) | 3 (30%) | 0 (0%) | 126 |
| 2 | 116 (43.4%) | 10 (66.7%) | 5 (50%) | 15 (100%) | 146 |
| Total | 267 | 15 | 10 | 15 | 307 |

$\chi^2 = 22.389$ ; $p = 0.001$

**Table 3.** Co-expression of IGF-1R-EHD1 - Frequencies considering all tissue types and only primary tumors:<colcnt=3>

| IGF-1R-EHD1 |  |  |
| --- | --- | --- |
|  | All tissues | Primary tumors |
| Negative | 4 (1.9%) | 4 (2.2%) |
| EHD1 or IGF-1R positive | 84 (39.1%) | 76 (41.5%) |
| Both positive | 127 (59.1%) | 103 (56.3%) |
| Total | 183 | 183 |

**Table 4.**
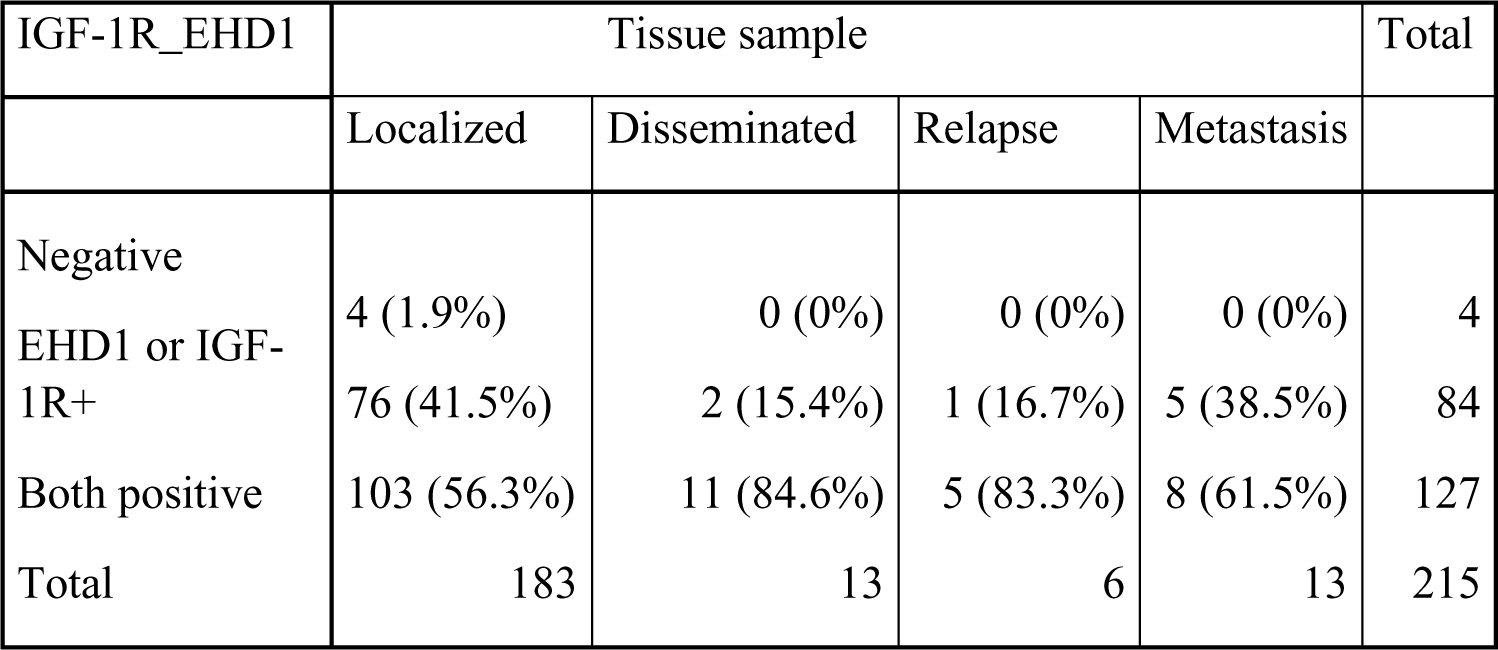
Correlation between IGF-1R-EHD1 co-expression and tissue types:<colcnt=6>

| IGF-1R_EHD1 | Tissue sample |  |  |  | Total |
| --- | --- | --- | --- | --- | --- |
|  | Localized | Disseminated | Relapse | Metastasis |  |
| Negative | 4 (1.9%) | 0 (0%) | 0 (0%) | 0 (0%) | 4 |
| EHD1 or IGF-1R+ | 76 (41.5%) | 2 (15.4%) | 1 (16.7%) | 5 (38.5%) | 84 |
| Both positive | 103 (56.3%) | 11 (84.6%) | 5 (83.3%) | 8 (61.5%) | 127 |
| Total | 183 | 13 | 6 | 13 | 215 |

**Table 5.**
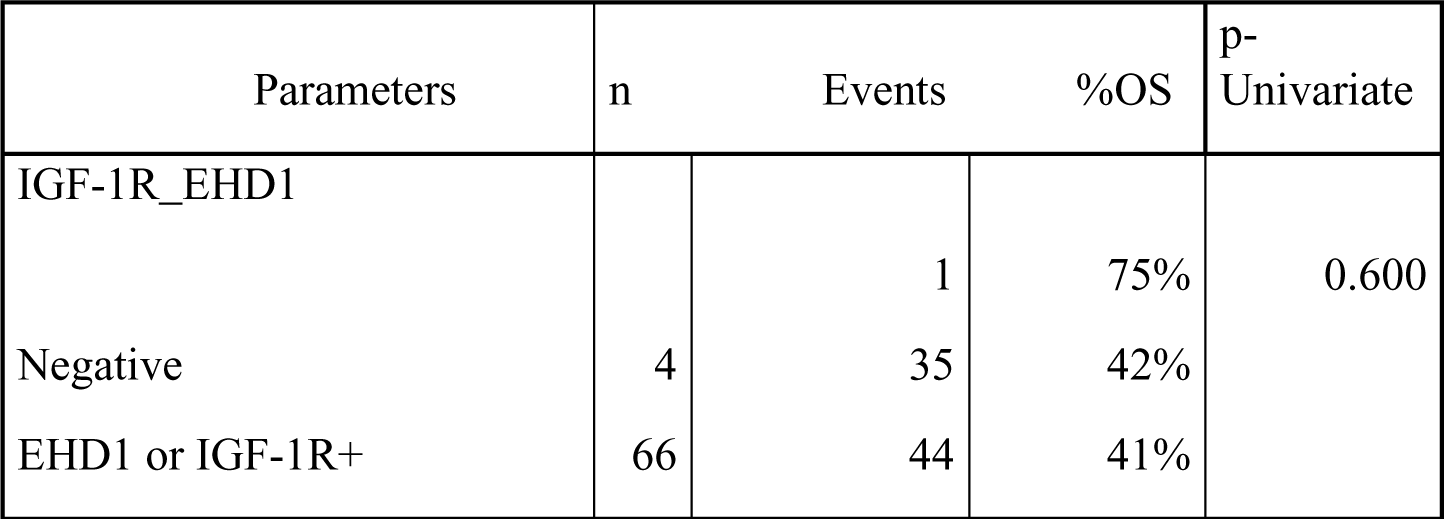
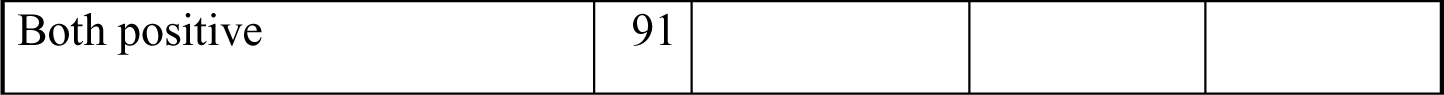
Association between IGF-1R-EHD1 co-expression and Overall survival (OS):

**Table 6.** Association between IGF-1R-EHD1 co-expression and Progression free survival (PFS):

| Parameters | n | Events | %PFS | p-Univariate |
| --- | --- | --- | --- | --- |
| IGF-1R_EHD1 |  |  |  |  |
| Negative | 4 | 1 | 75% | 0.334 |
| EHD1 or IGF-1R + | 66 | 38 | 34.4% |  |
| Both positive | 92 | 43 | 47% |  |

**Table 7.**
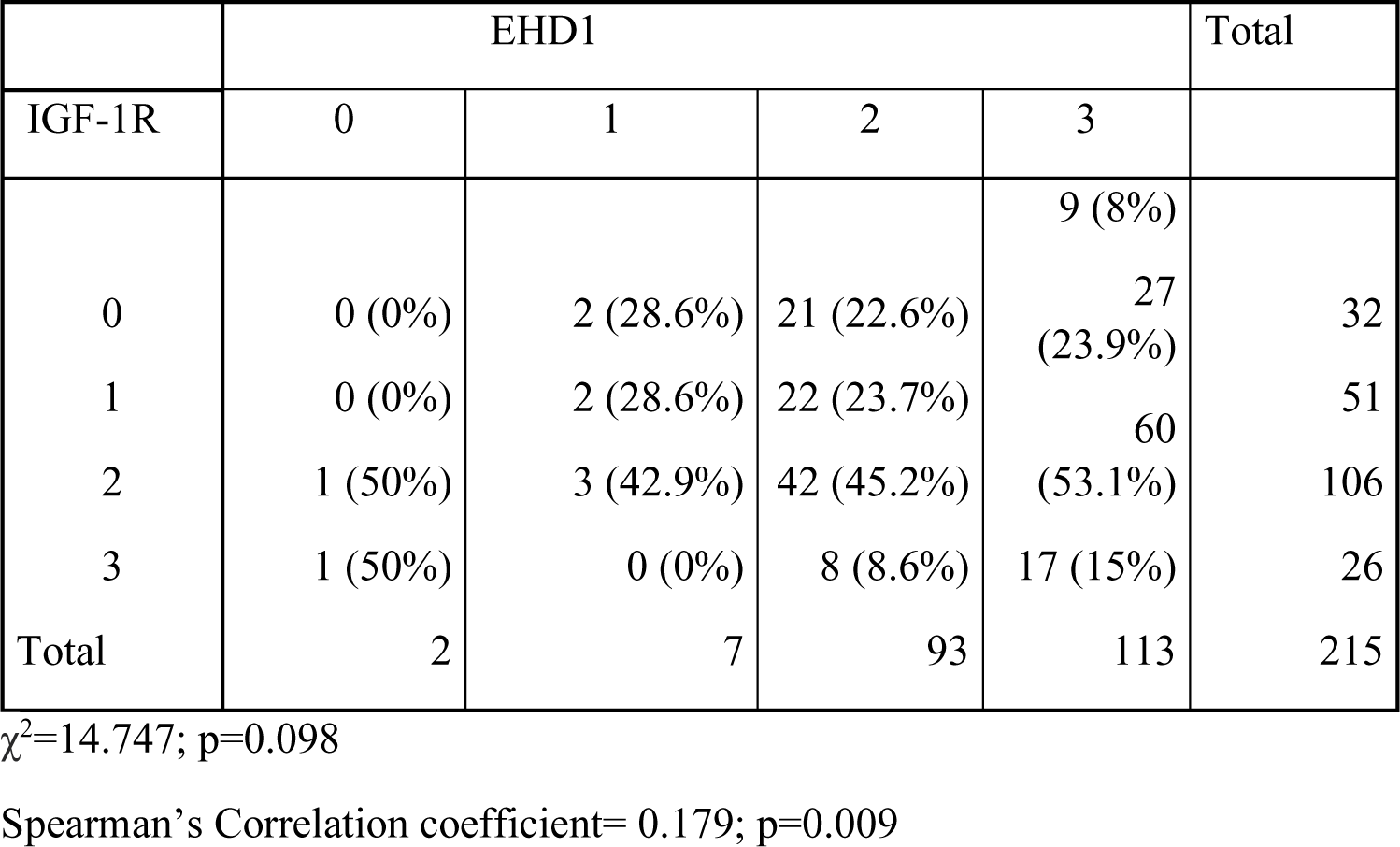
Correlation between IGF-1R and EHD1 IHC expression (4 categories):

## REFERENCES

1. Naslavsky N, Caplan S. EHD proteins: key conductors of endocytic transport. Trends Cell Biol 21, 122–131 (2011).

2. George M, et al. Shared as well as distinct roles of EHD proteins revealed by biochemical and functional comparisons in mammalian cells and C. elegans. BMC Cell Biol 8, 3 (2007).

3. Daumke O, Lundmark R, Vallis Y, Martens S, Butler PJ, McMahon HT. Architectural and mechanistic insights into an EHD ATPase involved in membrane remodelling. Nature 449, 923–927 (2007).

4. Cypher LR, et al. CSF-1 receptor signalling is governed by pre-requisite EHD1 mediated receptor display on the macrophage cell surface. Cell Signal 28, 1325–1335 (2016).

5. Tom EC, et al. EHD1 and RUSC2 Control Basal Epidermal Growth Factor Receptor Cell Surface Expression and Recycling. Mol Cell Biol 40, (2020).

6. Lu H, Meng Q, Wen Y, Hu J, Zhao Y, Cai L. Increased EHD1 in non-small cell lung cancer predicts poor survival. Thorac Cancer 4, 422–432 (2013).

7. Gao Y, Wang Y, Sun L, Meng Q, Cai L, Dong X. Expression of TGFβ-1 and EHD1 correlated with survival of non-small cell lung cancer. Tumour Biol 35, 9371–9380 (2014).

8. Meng Q, et al. Mammalian Eps15 homology domain 1 promotes metastasis in non-small cell lung cancer by inducing epithelial-mesenchymal transition. Oncotarget 8, 22433–22442 (2017).

9. Gao J, Meng Q, Zhao Y, Chen X, Cai L. EHD1 confers resistance to cisplatin in non-small cell lung cancer by regulating intracellular cisplatin concentrations. BMC Cancer 16, 470 (2016).

10. Tong D, et al. Increased Eps15 homology domain 1 and RAB11FIP3 expression regulate breast cancer progression via promoting epithelial growth factor receptor recycling. Tumour Biol 39, 1010428317691010 (2017).

11. Wang X, et al. NF-κB-driven improvement of EHD1 contributes to erlotinib resistance in EGFR-mutant lung cancers. Cell Death Dis 9, 418–418 (2018).

12. Liu Y, et al. Eps15 homology domain 1 promotes the evolution of papillary thyroid cancer by regulating endocytotic recycling of epidermal growth factor receptor. Oncol Lett 16, 4263–4270 (2018).

13. Wang T, et al. Mammalian Eps15 homology domain 1 potentiates angiogenesis of non-small cell lung cancer by regulating β2AR signaling. J Exp Clin Cancer Res 38, 174 (2019).

14. Huang J, et al. A feedback circuit comprising EHD1 and 14-3-3ζ sustains β-catenin/c-Myc-mediated aerobic glycolysis and proliferation in non-small cell lung cancer. Cancer Lett 520, 12–25 (2021).

15. Lu Y, Wang W, Tan S. EHD1 promotes the cancer stem cell (CSC)-like traits of glioma cells via interacting with CD44 and suppressing CD44 degradation. Environ Toxicol 37, 2259–2268 (2022).

16. Huang J, et al. Targeting the IL-1β/EHD1/TUBB3 axis overcomes resistance to EGFR-TKI in NSCLC. Oncogene 39, 1739-1755 (2020).

17. Jin W. The Role of Tyrosine Kinases as a Critical Prognostic Parameter and Its Targeted Therapies in Ewing Sarcoma. Frontiers in Cell and Developmental Biology 8, (2020).

18. Abella JV, Park M. Breakdown of endocytosis in the oncogenic activation of receptor tyrosine kinases. Am J Physiol Endocrinol Metab 296, E973–984 (2009).

19. Di Fiore PP, De Camilli P. Endocytosis and signaling. an inseparable partnership. Cell 106, 1–4 (2001).

20. Sigismund S, Confalonieri S, Ciliberto A, Polo S, Scita G, Di Fiore PP. Endocytosis and signaling: cell logistics shape the eukaryotic cell plan. Physiol Rev 92, 273–366 (2012).

21. Mosesson Y, Mills GB, Yarden Y. Derailed endocytosis: an emerging feature of cancer. Nat Rev Cancer 8, 835–850 (2008).

22. Grier HE. The Ewing family of tumors. Ewing’s sarcoma and primitive neuroectodermal tumors. Pediatr Clin North Am 44, 991-1004 (1997).

23. Vornicova O, Bar-Sela G. Investigational therapies for Ewing sarcoma: a search without a clear finding. Expert Opinion on Investigational Drugs 25, 679–686 (2016).

24. Ginsberg JP, et al. EWS-FLI1 and EWS-ERG gene fusions are associated with similar clinical phenotypes in Ewing’s sarcoma. J Clin Oncol 17, (1999).

25. Miser JS, et al. Treatment of metastatic Ewing’s sarcoma or primitive neuroectodermal tumor of bone: evaluation of combination ifosfamide and etoposide--a Children’s Cancer Group and Pediatric Oncology Group study. J Clin Oncol 22, (2004).

26. Siligan C, et al. EWS-FLI1 target genes recovered from Ewing’s sarcoma chromatin. Oncogene 24, (2005).

27. Herrero-Martin D, et al. Stable interference of EWS-FLI1 in an Ewing sarcoma cell line impairs IGF-1/IGF-1R signalling and reveals TOPK as a new target. Br J Cancer 101, (2009).

28. Toretsky JA, Kalebic T, Blakesley V, LeRoith D, Helman LJ. The insulin-like growth factor-I receptor is required for EWS/FLI-1 transformation of fibroblasts. J Biol Chem 272, 30822–30827 (1997).

29. Cironi L, et al. IGF1 is a common target gene of Ewing’s sarcoma fusion proteins in mesenchymal progenitor cells. PLoS One 3, e2634 (2008).

30. Prieur A, Tirode F, Cohen P, Delattre O. EWS/FLI-1 silencing and gene profiling of Ewing cells reveal downstream oncogenic pathways and a crucial role for repression of insulin-like growth factor binding protein 3. Mol Cell Biol 24, 7275–7283 (2004).

31. Osher E, Macaulay VM. Therapeutic Targeting of the IGF Axis. Cells 8, (2019).

32. Brahmkhatri VP, Prasanna C, Atreya HS. Insulin-like growth factor system in cancer: novel targeted therapies. Biomed Res Int 2015, 538019 (2015).

33. Christopoulos PF, Msaouel P, Koutsilieris M. The role of the insulin-like growth factor-1 system in breast cancer. Mol Cancer 14, 43 (2015).

34. Arcaro A. Targeting the insulin-like growth factor-1 receptor in human cancer. Front Pharmacol 4, 30 (2013).

35. Scotlandi K, et al. Insulin-like growth factor I receptor-mediated circuit in Ewing’s sarcoma/peripheral neuroectodermal tumor: a possible therapeutic target. Cancer Res 56, 4570–4574 (1996).

36. Juergens H, et al. Preliminary efficacy of the anti-insulin-like growth factor type 1 receptor antibody figitumumab in patients with refractory Ewing sarcoma. J Clin Oncol 29, 4534–4540 (2011).

37. Anderson PM, et al. A phase II study of clinical activity of SCH 717454 (robatumumab) in patients with relapsed osteosarcoma and Ewing sarcoma. Pediatr Blood Cancer 63, 1761–1770 (2016).

38. Tolcher AW, et al. Phase I, pharmacokinetic, and pharmacodynamic study of AMG 479, a fully human monoclonal antibody to insulin-like growth factor receptor 1. J Clin Oncol 27, 5800–5807 (2009).

39. Olmos D, et al. Safety, pharmacokinetics, and preliminary activity of the anti-IGF-1R antibody figitumumab (CP-751,871) in patients with sarcoma and Ewing’s sarcoma: a phase 1 expansion cohort study. Lancet Oncol 11, 129–135 (2010).

40. Kurzrock R, et al. A phase I study of weekly R1507, a human monoclonal antibody insulin-like growth factor-I receptor antagonist, in patients with advanced solid tumors. Clin Cancer Res 16, 2458–2465 (2010).

41. Malempati S, et al. Phase I/II trial and pharmacokinetic study of cixutumumab in pediatric patients with refractory solid tumors and Ewing sarcoma: a report from the Children’s Oncology Group. J Clin Oncol 30, 256–262 (2012).

42. Murakami H, et al. Phase 1 study of ganitumab (AMG 479), a fully human monoclonal antibody against the insulin-like growth factor receptor type I (IGF1R), in Japanese patients with advanced solid tumors. Cancer Chemother Pharmacol 70, 407–414 (2012).

43. Schoffski P, et al. An open-label, phase 2 study evaluating the efficacy and safety of the anti-IGF-1R antibody cixutumumab in patients with previously treated advanced or metastatic soft-tissue sarcoma or Ewing family of tumours. Eur J Cancer 49, 3219–3228 (2013).

44. Qu X, et al. Update of IGF-1 receptor inhibitor (ganitumab, dalotuzumab, cixutumumab, teprotumumab and figitumumab) effects on cancer therapy. Oncotarget 8, 29501–29518 (2017).

45. Baserga R. The decline and fall of the IGF-I receptor. J Cell Physiol 228, 675–679 (2013).

46. Ebinger S, et al. Characterization of Rare, Dormant, and Therapy-Resistant Cells in Acute Lymphoblastic Leukemia. Cancer Cell 30, 849–862 (2016).

47. Naslavsky N, Rahajeng J, Sharma M, Jovic M, Caplan S. Interactions between EHD proteins and Rab11-FIP2: a role for EHD3 in early endosomal transport. Mol Biol Cell 17, 163–177 (2006).

48. Rotem-Yehudar R, Galperin E, Horowitz M. Association of insulin-like growth factor 1 receptor with EHD1 and SNAP29. J Biol Chem 276, 33054–33060 (2001).

49. Crudden C, et al. Below the Surface: IGF-1R Therapeutic Targeting and Its Endocytic Journey. Cells 8, (2019).

50. Fernando R, Caldera O, Smith TJ. Therapeutic IGF-I receptor inhibition alters fibrocyte immune phenotype in thyroid-associated ophthalmopathy. Proc Natl Acad Sci U S A 118, (2021).

51. Li SL, Kato J, Paz IB, Kasuya J, Fujita-Yamaguchi Y. Two new monoclonal antibodies against the alpha subunit of the human insulin-like growth factor-I receptor. Biochem Biophys Res Commun 196, 92–98 (1993).

52. Suva LJ, Washam C, Nicholas RW, Griffin RJ. Bone metastasis: mechanisms and therapeutic opportunities. Nat Rev Endocrinol 7, 208–218 (2011).

53. Jović M, Naslavsky N, Rapaport D, Horowitz M, Caplan S. EHD1 regulates beta1 integrin endosomal transport: effects on focal adhesions, cell spreading and migration. J Cell Sci 120, 802–814 (2007).

54. Iseka FM, et al. Role of the EHD Family of Endocytic Recycling Regulators for TCR Recycling and T Cell Function. J Immunol 200, 483–499 (2018).

55. Foti M, Moukil MA, Dudognon P, Carpentier JL. Insulin and IGF-1 receptor trafficking and signalling. Novartis Found Symp 262, 125–141; discussion 141-127, 265-128 (2004).

56. de Groot S, Röttgering B, Gelderblom H, Pijl H, Szuhai K, Kroep JR. Unraveling the Resistance of IGF-Pathway Inhibition in Ewing Sarcoma. Cancers (Basel) 12, (2020).

57. Rieger L, O’Connor R. Controlled Signaling-Insulin-Like Growth Factor Receptor Endocytosis and Presence at Intracellular Compartments. Front Endocrinol (Lausanne) 11, 620013 (2020).

58. Romanelli RJ, LeBeau AP, Fulmer CG, Lazzarino DA, Hochberg A, Wood TL. Insulin-like growth factor type-I receptor internalization and recycling mediate the sustained phosphorylation of Akt. J Biol Chem 282, 22513–22524 (2007).

59. Demonbreun AR, et al. Myoferlin is required for insulin-like growth factor response and muscle growth. Faseb j 24, 1284–1295 (2010).

60. Essandoh K, et al. Tsg101 positively regulates physiologic-like cardiac hypertrophy through FIP3-mediated endosomal recycling of IGF-1R. Faseb j 33, 7451–7466 (2019).

61. Chen G, et al. GIGYF1 disruption associates with autism and impaired IGF-1R signaling. J Clin Invest 132, (2022).

62. Rieger L, O’Shea S, Godsmark G, Stanicka J, Kelly G, O’Connor R. IGF-1 receptor activity in the Golgi of migratory cancer cells depends on adhesion-dependent phosphorylation of Tyr(1250) and Tyr(1251). Sci Signal 13, (2020).

63. Agarwal NK, et al. Smoothened (SMO) regulates insulin-like growth factor 1 receptor (IGF1R) levels and protein kinase B (AKT) localization and signaling. Lab Invest 102, 401–410 (2022).

64. Bhattacharyya S, et al. Endocytic recycling protein EHD1 regulates primary cilia morphogenesis and SHH signaling during neural tube development. Sci Rep 6, 20727 (2016).

65. Gazdar AF. Activating and resistance mutations of EGFR in non-small-cell lung cancer: role in clinical response to EGFR tyrosine kinase inhibitors. Oncogene 28, S24–S31 (2009)..

66. Liu Y, et al. A novel EHD1/CD44/Hippo/SP1 positive feedback loop potentiates stemness and metastasis in lung adenocarcinoma. Clin Transl Med 12, e836 (2022).

67. López-Guerrero JA, et al. Clinicopathological significance of cell cycle regulation markers in a large series of genetically confirmed Ewing’s sarcoma family of tumors. Int J Cancer 128, 1139–1150 (2011).

68. Kim D, Paggi JM, Park C, Bennett C, Salzberg SL. Graph-based genome alignment and genotyping with HISAT2 and HISAT-genotype. Nat Biotechnol 37, 907–915 (2019).

69. Kovaka S, Zimin AV, Pertea GM, Razaghi R, Salzberg SL, Pertea M. Transcriptome assembly from long-read RNA-seq alignments with StringTie2. Genome Biol 20, 278 (2019).

70. Love MI, Huber W, Anders S. Moderated estimation of fold change and dispersion for RNA-seq data with DESeq2. Genome Biol 15, 550 (2014).

